# DNA damage drives a unique, Alzheimer’s disease-relevant senescent state in neurons

**DOI:** 10.64898/2026.04.02.716205

**Authors:** Jun-Wei B. Hughes, Anja Sandholm, Duncan Croll, Fiona Senchyna, Kevin Schneider, Rachel Butterfield, Tyne L M McHugh, Ian Brown, Hideto Deguchi, Tyler A. U. Hilsabeck, Sally Mak, Kenneth A. Wilson, Hayk Davtyan, Mathew Blurton-Jones, Joseph Herdy, Ryo Higuchi-Sanabria, Fred H. Gage, David Furman, Lisa M. Ellerby, Pierre-Yves Desprez, Judith Campisi

**Affiliations:** Buck Institute for Research on Aging, Novato, CA, USA; Leonard Davis School of Gerontology, University of Southern California, Los Angeles, CA, USA; Department of Neurobiology and Behavior, University of California, Irvine, CA, USA; The Salk Institute for Biological Studies, La Jolla, CA, USA; California Pacific Medical Center, Research Institute, San Francisco, California, USA

**Keywords:** Senescence, Alzheimer’s disease, DNA damage, neurons, weighted gene co-expression network analysis

## Abstract

Alzheimer’s disease (AD) shares molecular hallmarks with the canonical drivers of cellular senescence. Senescent cells have also been shown to accumulate in the brain with age, yet the mechanisms linking AD pathology to the accumulation of senescent cells in the brain remain unclear. Here, we demonstrate that DNA damage in patient-derived directly induced neurons (iNs) drives a senescent-like cell state with relevance to AD. DNA damage-induced senescent iNs show significant transcriptional concordance with human AD neurons and a weighted gene co-expression network analysis (WGCNA) uncovers candidate regulators associated with the senescent-like state in neurons. Direct comparison of iNs to the original patient fibroblasts reveals striking cell-type specific senescence signatures following DNA damage. iNs adopt a p21-associated senescent-like state characterized by a senescence-associated secretory phenotype (SASP) and predicted activation of NF-κꞴ1. In contrast, fibroblasts develop a p16-associated senescent state lacking a SASP phenotype and show a predicted repression of NF-κꞴ1. Early responses to DNA damage further reveal divergent DNA damage response (DDR), with neurons exhibiting higher accumulation of damage lesions relative to fibroblasts. Together, these findings demonstrate that DNA damage drives a unique senescent-like neuronal state that models molecular features of AD, while also revealing fundamental cell-type specific differences in senescent-like phenotypes and DDR.

## INTRODUCTION

Cellular senescence is canonically defined by stable cell cycle arrest, resistance to apoptosis, and the development of a senescence-associated secretory phenotype (SASP), a pro-inflammatory secretory program characterized by the release of cytokines, chemokines, growth factors, and proteases^1,2^. Because neurons are post-mitotic, they were previously thought to be incapable of undergoing senescence. In addition, neurons are commonly considered less secretory and inflammatory than other neural cells in the brain, including glial cells^3–5^. However, emerging evidence challenges these assumptions and shows that neurons can adopt a senescent-like state across many aging and disease contexts, including an inflammatory signature like the canonical SASP^6–12^.

Accumulating studies identify a senescent-like state in neurons in various Alzheimer’s Disease (AD) contexts^6–12^. AD is the most common form of dementia and is characterized by progressive neurodegeneration, synaptic loss, and the accumulation of neuropathologic hallmarks, including amyloid-beta plaques and neurofibrillary tangles^13–17^. Despite growing evidence for the presence of senescent cells in AD brains, the mechanisms driving their accumulation and contribution to AD pathology remain poorly understood.

CDKN2A/p16 and CDKN1A/p21 are key regulators of cell cycle arrest after senescence induction, usually by p21 initiating the early senescence response and p16 maintaining the later senescence response, but it is still a major question how these two cell cycle inhibitors are engaged in cell types that are already cell-cycle arrested^1,18^. The first study to address this question in neurons revealed senescent-like neurons in older mice with a p21-dependent phenotype that was linked to the DNA damage response (DDR)^6^. More recent studies built on this finding and showed a p21-associated senescence response in older rat brain neurons and in neurons derived from induced-pluripotent stem cells (iPSC) in the context of Parkinson’s disease (PD)^7,10^. As p16 usually maintains a senescent cell state, studies have also shown p16-associated senescence phenotypes in neural stem cells (NSCs) and medium spiny neurons (MSNs) from iPSCs in the context of Huntington’s disease (HD)^8^. Additionally, patient-derived directly-induced neurons (iNs) from AD patients have shown a p16-associated senescence response in the AD iNs^9^. Multiple studies have identified p16 as being elevated in post-mortem human AD brains as well^9,19^. Interestingly, a single-cell RNA-sequencing (scRNA-seq) study from human AD brains highlighted a CDKN2D/p19-expressing neuron population with senescent-like phenotypes^11^. Altogether, p16 and p21 likely both play a role in neuronal senescence with aging and neurodegenerative diseases but the cause of the accumulation of these senescent neurons is unknown.

DNA damage is widely implicated in senescent phenotypes, including the senescent neuron phenotype, but the direct link of genotoxic stress driving a senescent state in human neurons has yet to be made. This is also important in the context of AD as several major molecular hallmarks of AD, such as DNA damage increase in AD, overlap with known inducers of cellular senescence, such as DNA-damage induced senescence^1,14^. As DNA damage is thought to be a primary driver of senescence *in vivo*, we hypothesized that DNA damage-induced senescence drives AD-relevant neuronal phenotypes^1,14^. To test this, we use the iN model, which retains age-associated transcriptomic and epigenetic features that are largely reset in iPSC models^12^. iNs are derived from primary, dermal fibroblasts and induced into neurons through ASCL1 and NGN2 overexpression, two pro-neuronal, pioneer transcription factors (TFs), skipping the stem cell state and differentiating into adult neurons within 3 weeks^12^.

Using the iN model, we demonstrate that DNA-damage induces a senescent-like state in iNs that models key features of AD neurons. DNA-damaged iNs showed strong concordance with AD iNs, including similarities with scRNA-seq datasets from post-mortem human AD brains. Weighted gene co-expression network analysis (WGCNA) identified candidate regulators associated with senescent-like state in iNs. To determine whether these phenotypes were unique to neurons, we directly compared DNA-damaged iNs to the donor-matched fibroblasts from which they were derived. These analyses revealed striking cell-type specific senescence programs. DNA-damaged iNs exhibited a pronounced SASP profile relative to fibroblasts, likely driven by predicted differential activation of NF-κꞴ1 between cell states. Consistent with these findings, we identified a p21-associated senescence response in iNs while fibroblasts exhibited a p16-associated senescence response. Early DDR also differed between cell types, ultimately leading to higher damage signaling in neurons.

## RESULTS

### DNA damaged iNs and AD neurons share synaptic and inflammatory gene expression dysregulation

Given the overlap between major molecular hallmarks of AD and canonical inducers of senescence, combined with the evidence that DNA damage is a primary driver of senescence *in vivo*, we exposed iNs from 9 healthy donors to 15 Grays of γ-irradiation (IR) to induce a senescence-like phenotype in iNs (Fig. 1A-C). 15 Grays of IR is a well-established method that triggers senescence primarily through DNA double-strand breaks (DSBs)^2,20–22^. Bulk RNA-sequencing (RNA-seq) was performed 14 days after IR and all analyses were performed relative to donor-matched non-irradiated controls to isolate senescence-specific responses (Fig. 1C). We found that iNs exposed to IR upregulated inflammatory and senescence-associated genes and pathways, such as CDKN1A and p53-related signaling, both well-known modulators of many senescence contexts (Fig. 1D-E). Notably, KEGG enrichment of the IR iN upregulated genes also predicted that the NF-κꞴ signaling pathway was activated, which is an important regulator of the SASP (Supp. Fig. 1)^23^. Irradiated iNs also displayed a significant downregulation of synapse-associated genes, suggesting synapse dysregulation after DNA damage (Fig. 1E).

**Figure 1.**
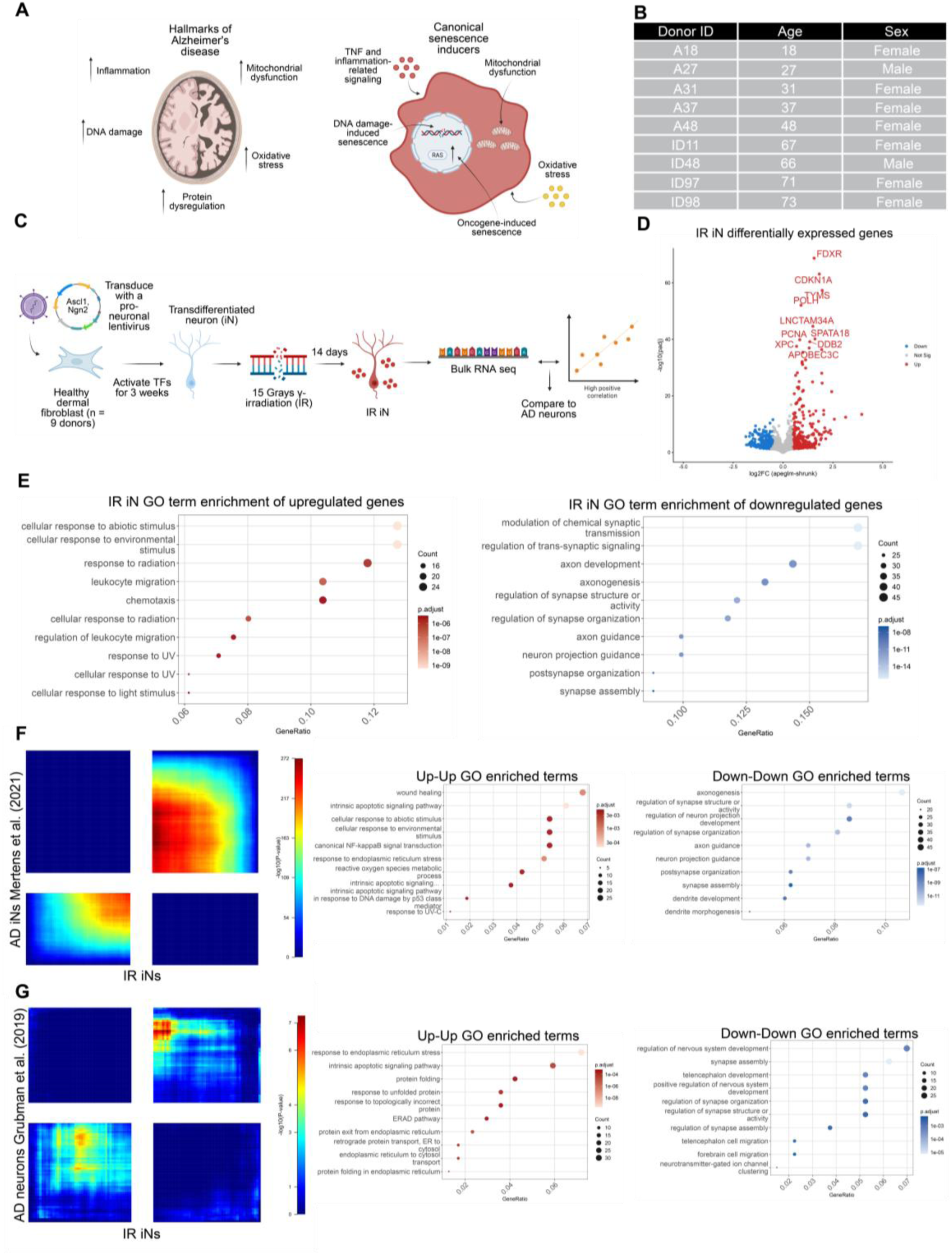
DNA-damaged iNs and AD neurons share synaptic and inflammatory gene expression dysregulation (A) Schematic of molecular hallmarks of AD and canonical inducers of cellular senescence. (B) Table of 9 healthy donor fibroblasts used for direct differentiation experiments in all of the experiments with donor ID, age, and sex provided. (C) Schematic of transdifferentiation system and DNA damage-induced senescence. 9 healthy dermal fibroblasts between ages 18-73 were transdifferentiated into iNs and induced to senesce by γ-irradiation (IR). RNA was collected for bulk RNA sequencing (RNA-seq) 14 days post IR from 1-2 technical replicates per donor in each condition. (D) Volcano plot of differentially expressed genes (DEGs) in healthy IR vs CTL iNs. DEG cutoffs were defined by |log2FC| > 0.5 (log2foldchange) and adjusted p-value < 0.05. The top 10 DEGs by significance are labeled and the x-axis is limited to a log2FC range of −5 to 5. (E) Top 10 GO enrichment terms from upregulated (left) and downregulated (right) DEGs from the IR iNs. (F-G) RRHO of IR iN DEGs compared to AD iNs from Mertens et al. (2021) (F) or pseudobulked neurons from Grubman et al. (2019) (G). Top 10 GO enrichment terms provided for concordant quadrants in each comparative study. The bottom left quadrant of the RRHO indicates concordant upregulated genes between datasets and the top right quadrant of the RRHO indicates concordant downregulated genes between datasets.

To determine if the senescent-like state induced by IR in iNs was relevant to AD, we used rank-rank hypergeometric overlap (RRHO) to compare our IR iN dataset with previously published AD iN datasets and scRNA-seq datasets from AD patients^24^. RRHO was used because it enables a threshold-free, genome-wide comparison of two ranked gene lists by applying a hypergeometric test across all possible rank cutoffs simultaneously^24^. This avoids the arbitrary imposition of significance or fold-change thresholds, which can exclude biologically relevant genes, and allows detection of both concordant and discordant transcriptomic patterns across the full expression spectrum^24^. RRHO analysis between IR iNs and AD iNs showed significant overlap between up and down regulated genes (Fig. 1F)^12^. Specifically, inflammation-related signaling increased in expression and synapse-associated signaling decreased in expression in both datasets, suggesting iNs exposed to IR may exhibit similar transcriptomic profiles as AD iNs (Fig. 1F).

To determine whether the transcriptional changes observed following IR exposure in iNs were also present in human AD brains, we compared our dataset with two previously published AD brain scRNA-seq datasets. The first dataset consisted of neurons from the entorhinal cortex of 6 individuals with Braak-characterized AD diagnosis, while the second dataset was derived from the prefrontal cortex neurons of 427 individuals from the “The Religious Orders Study and Memory and Aging Project” (ROSMAP), which characterizes AD progression using Braak staging, Cerad score, and cognitive testing (Cogdx)^25,26^. Braak staging and Cerad scoring are used to measure pathologic burden in AD brains while Cogdx scoring is used to measure cognitive impairment in AD patients^16,17,26^. Both scRNA-seq datasets were pseudobulked based on neurons prior to RRHO analysis, and the ROSMAP dataset was more specifically pseudobulked based on excitatory neurons, to ensure proper neuron-to-neuron comparisons across datasets.

RRHO analysis between IR iNs and entorhinal cortex AD neurons showed moderate overlap, with shared upregulated genes enriched with endoplasmic reticulum and lysosomal stress and shared downregulated genes associated with synapse dysregulation (Fig. 1G). In contrast, comparison of IR iNs to the ROSMAP AD excitatory neurons revealed discordant signatures across AD classifications based on Cerad score, Braak stage, or Cogdx (Supp. Fig. 2). Upregulated genes in the ROSMAP AD excitatory neurons were related to synapse-associated signaling, which was downregulated in IR iNs, whereas downregulated genes in the ROSMAP AD excitatory neurons were related to metabolic signaling, which was upregulated in IR iNs (Supp. Fig. 2). Together, these results indicate that DNA damage-induced senescence in iNs models key transcriptional features in AD iNs, while showing both concordant and discordant gene expression signatures relative to scRNA-seq datasets from human AD brains.

**Figure 2.**
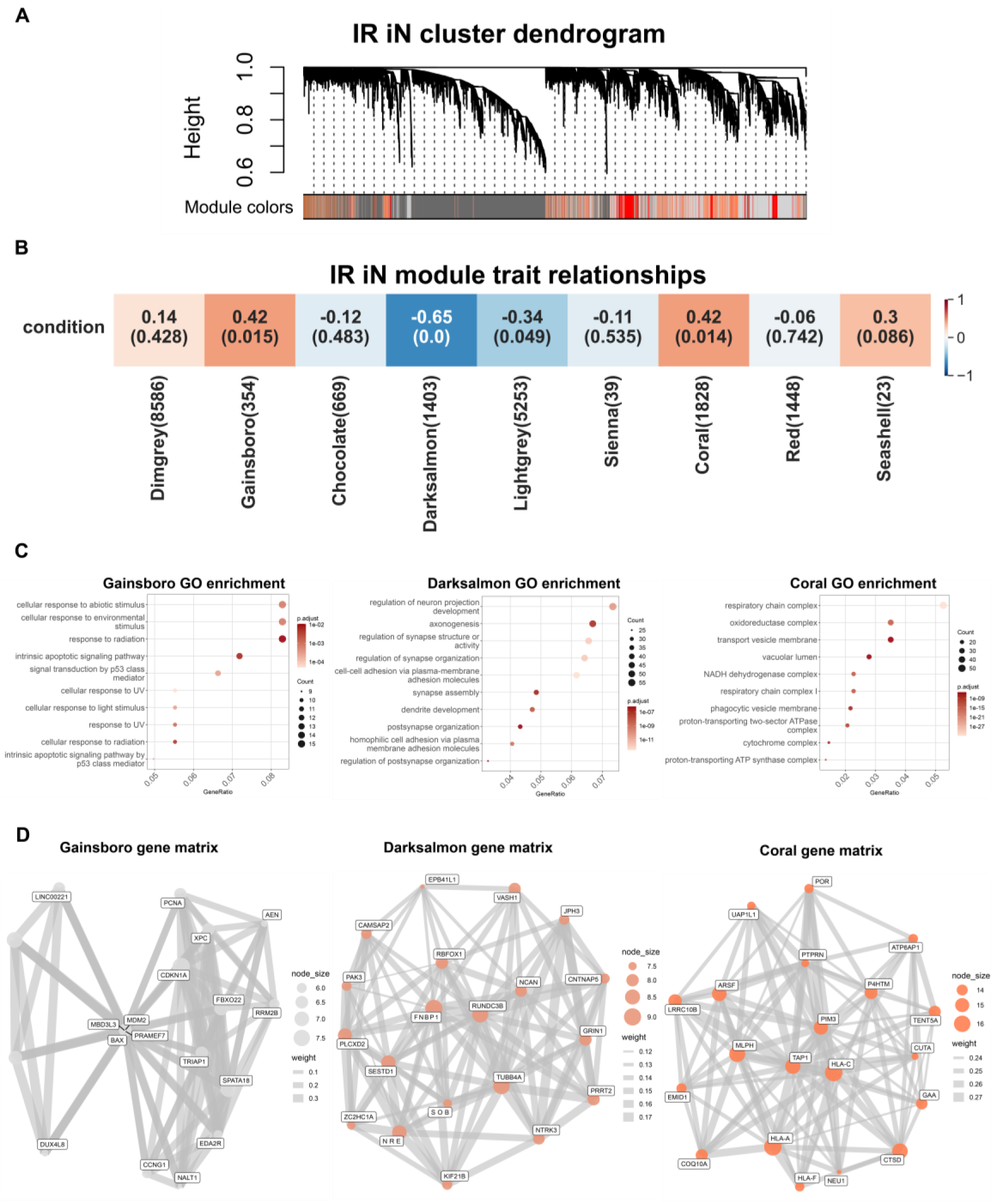
Weighted gene co-expression network analysis reveals candidate drivers of the IR iN cell state (A) Hierarchical clustering dendrogram of DEGs based on dissimilarity of the topological overlap from the WGCNA. The color bar below the dendrogram indicates module assignments, with each color representing a distinct co-expression module of highly correlated genes. (B) WGCNA module-trait heatmap relationships from Fig. 1 RNA-seq input of 9 healthy iNs after IR. Our iN sample number gave us confidence that the WGCNA would reach sufficient power for meaningful results, as Langfelder and Horvath recommend using at 15 samples and our total sample number was 18 and 1-2 biological replicates per sample^27^. Rows represent IR condition, and columns represent module eigengenes. Each cell includes corresponding correlation to the specific module and correlation is indicated by the value in the box. Correlation ranges from -1 to 1 and blue indicates a negative correlation while red indicates a positive correlation. Correlation values are above the number in parenthesis which indicates the p-value of each module’s connectivity. (C) Top 10 GO enrichment terms of genes from the Gainsboro (left), Darksalmon (middle), and Coral (right) modules. (D) Module gene network visualization for the Gainsboro (left), Darksalmon (middle), and Coral (right) modules. Gene matrices were made using the BioMart R package. Nodes represent the top module genes ranked by connectivity z-score, node size scales with intramodular connectivity, and edges represent weighted topological overlap matrix (TOM) connections (TOM ≥ 0.001; top 8 connections retained per gene).

### Weighted gene co-expression network analysis reveals candidate drivers of the IR iN cell state

Having established that DNA damage-induced senescence in iNs models AD-relevant transcriptional phenotypes, we next sought to identify candidate genes that contribute to driving this senescence-like state in iNs. To do so, we performed a WGCNA to identify co-expression modules from the RNA-seq data used in Fig. 1 and uncover genes correlated with the IR condition in iNs, complementing our differential expression analysis with a network-based approach^27^.

WGCNA identified multiple highly connected and significantly correlated modules to the IR iN cell state, including the Gainsboro, Darksalmon, and Coral modules (Fig. 2B). Gene ontology (GO) enrichment analysis revealed that these modules corresponded to distinct biological mechanisms: the Gainsboro module was enriched for p53 signaling and inflammatory pathways, the Darksalmon module for synapse-associated signaling, and the Coral module for metabolism-related pathways (Fig. 2C). Notably, the Gainsboro and Darksalmon module genes enriched in GO terms similar to the GO terms from the RNA-seq data from IR iNs in Fig. 1E, reinforcing that these modules and pathways are important for the overall cell state. Within each module, highly connected genes, often referred to as hub genes, were identified based on connectivity z-score and visualized in network plots. CDKN1A emerged as one of the most highly connected hub genes in the Gainsboro module (Fig. 2D), which was also identified in the RNA-seq analysis as one of the highest differentially expressed genes (DEG) in IR iNs from Fig. 1D. These findings build on our differential expression analysis and suggest that the senescence-associated response to DNA damage in iNs may be driven by CDKN1A.

### Direct comparison of neurons to fibroblasts reveal distinct senescence-associated transcriptional signatures

A key advantage of the direct differentiation system is the ability to compare two distinct cell types from the same donor, thereby controlling for differences driven by donor-to-donor heterogeneity. To leverage this advantage, we exposed both fibroblasts and iNs, from the same donor, to 15 Grays IR and performed RNA-seq 14 days post-irradiation (Fig. 3A). Differential expression analysis of IR fibroblasts revealed robust transcriptional changes characteristic of canonical senescence programs. The most strongly upregulated genes were enriched for extracellular matrix (ECM) and cell adhesion pathways, while top downregulated genes were enriched for cell cycle progression, DNA replication, and DNA repair pathways, consistent with previously described senescence-associated transcriptional signatures (Fig. 3B-C)^28^.

**Figure 3.**
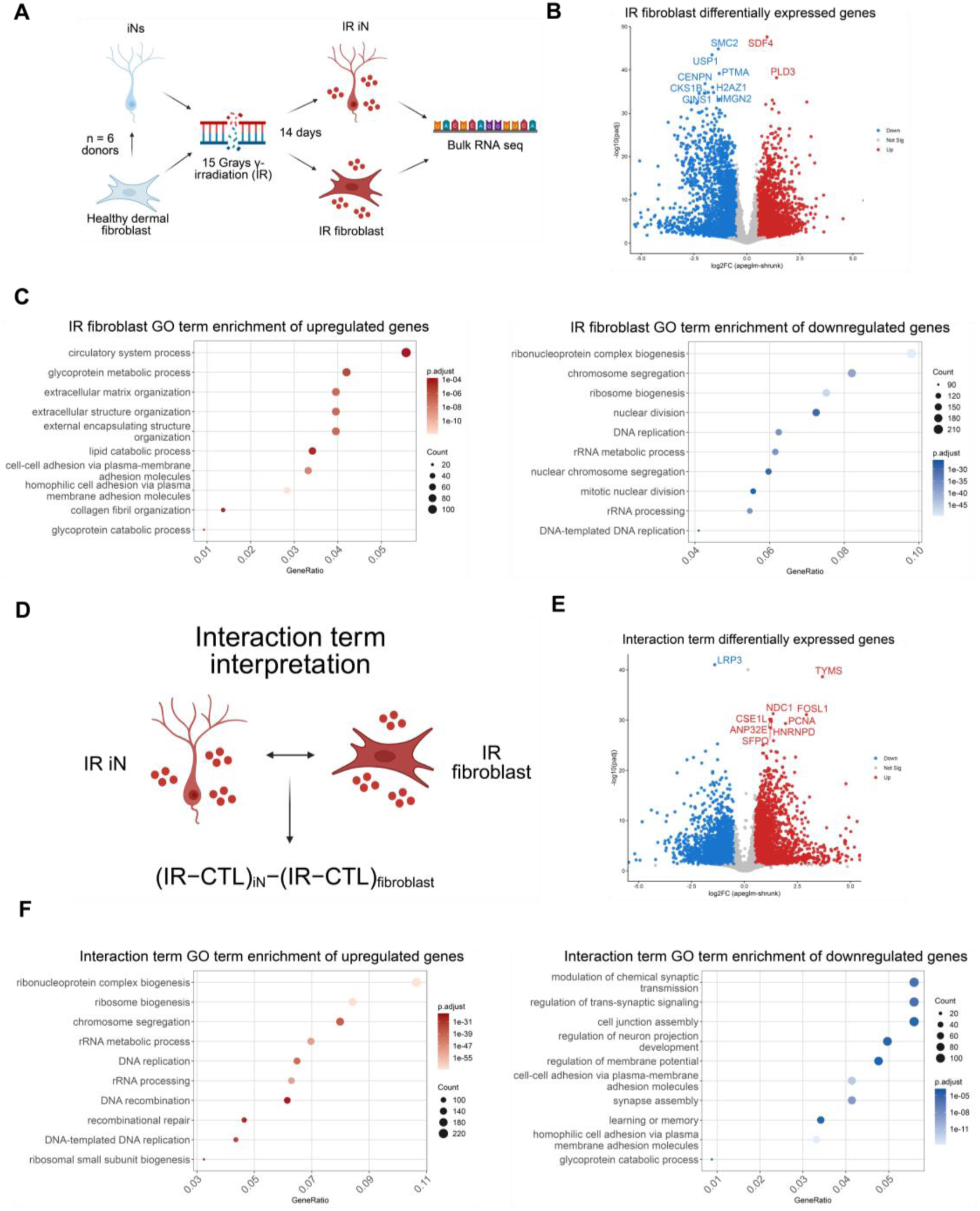
Direct comparison of neurons to fibroblasts reveal distinct senescence-associated transcriptional signatures (A) Schematic of DNA damage-induced senescence in iNs and fibroblasts from the same donor. 6 healthy dermal fibroblasts (donors: A18, A27, A31, A37, A48, ID97) were used for direct RNA-seq comparisons between cell types with 1-2 technical replicates per donor in each condition. (B) Volcano plot of differentially expressed genes (DEGs) in healthy IR vs CTL fibroblasts. DEG cutoffs were defined by |log2FC| > 0.5 and adjusted p-value < 0.05. The top 10 DEGs by significance are labeled and the x-axis is limited to a log2FC range of −5 to 5. (C) Top 10 GO enrichment terms from upregulated (left) and downregulated (right) DEGs from the IR fibroblasts. (D) Schematic illustrating the DESeq2 interaction term design, in which the effect of IR is first estimated within each cell type relative to its control, and then compared between iNs and fibroblasts to identify genes whose IR response differs by cell type. (E) Volcano plot of the interaction term. DEG cutoffs were defined by |log2FC| > 0.5 and adjusted p-value < 0.05. The top 10 DEGs by significance are labeled and the x-axis is limited to a log2FC range of −5 to 5. (F) Top 10 GO enrichment terms from upregulated (left) and downregulated (right) DEGs from the interaction term. Interaction term interpretations: Positive log2FC (interaction > 0) IR effect is more positive in iNs than in fibroblasts (either stronger up in iN, or less down in iN). Negative log2FC (interaction < 0) IR effect is more positive in fibroblasts than in iNs (either stronger up in fibroblasts, or stronger down in iN).

To directly compare senescence-associated responses between neurons and fibroblasts, we extracted DEGs from the interaction term of the DESeq2 model, enabling comparison of the IR response in iNs relative to fibroblasts while accounting for each cell type’s baseline expression (Fig. 3D). A positive value indicates the gene or pathway is either more highly expressed or less suppressed in iNs relative to fibroblasts, while a negative value indicates the converse (Fig. 3D). This analysis revealed striking cell-type specific transcriptional responses to DNA damage. Genes enriched in positive interaction terms were associated with cell cycle regulation and DNA replication and repair pathways, reflecting the stronger repression of these programs in IR-treated fibroblasts relative to iNs (Fig. 3E-F). In contrast, genes enriched in negative interaction terms were enriched in pathways associated with synapse signaling, showing a greater downregulation of synaptic programs in IR iNs relative to IR fibroblasts (Fig. 3E-F). Additionally, by comparing shared and unique DEGs between IR iNs and IR fibroblasts, we discovered only 32% of the IR iN upregulated DEGs and 14% of the IR iN downregulated DEGs were shared with the IR fibroblast DEGs (Supp. Fig. 3).

Together, these analyses demonstrate that DNA damage induces distinct senescence-associated transcriptional programs across cell types. While fibroblasts exhibit a canonical senescence response characterized by suppression of proliferation pathways, neurons display a distinct transcriptional program marked by synaptic dysfunction, suggesting tissue-specific effects and differences in cell cycle accessibility that drive divergent senescent fates.

### Senescence programs differ between neurons and fibroblasts after DNA damage and diverge from canonical senescence signatures

To determine whether the transcriptional programs induced by IR in iNs resemble previously described signatures of senescence or a distinct, neuron-specific program, we next performed gene set enrichment analysis (GSEA) using established senescence-associated gene sets, including CellAge and SenMayo, and applied to both the IR fibroblast and IR iN transcriptomic datasets (Fig. 4A)^22,29–35^. We found that both IR iN and IR fibroblast upregulated genes showed enrichment for the CellAge senescence signature, confirming activation of canonical senescence-associated transcriptional programs (Fig. 4B-C). Enrichment was notably weaker in the IR iNs for the CellAge senescence signature (Fig. 4B). Interestingly, IR iN upregulated genes also showed enrichment for the “SASP atlas fibroblast” signature, while IR fibroblasts downregulated genes were enriched for this signature (Fig. 4B-C). This unexpected pattern suggests that IR iNs may exhibit a more robust SASP than IR fibroblasts. Overall, these results indicate that DNA damage induces senescence-associated transcriptional programs in both neurons and fibroblasts, although novel cell type specific responses, particularly in the SASP, also exist.

**Figure 4.**
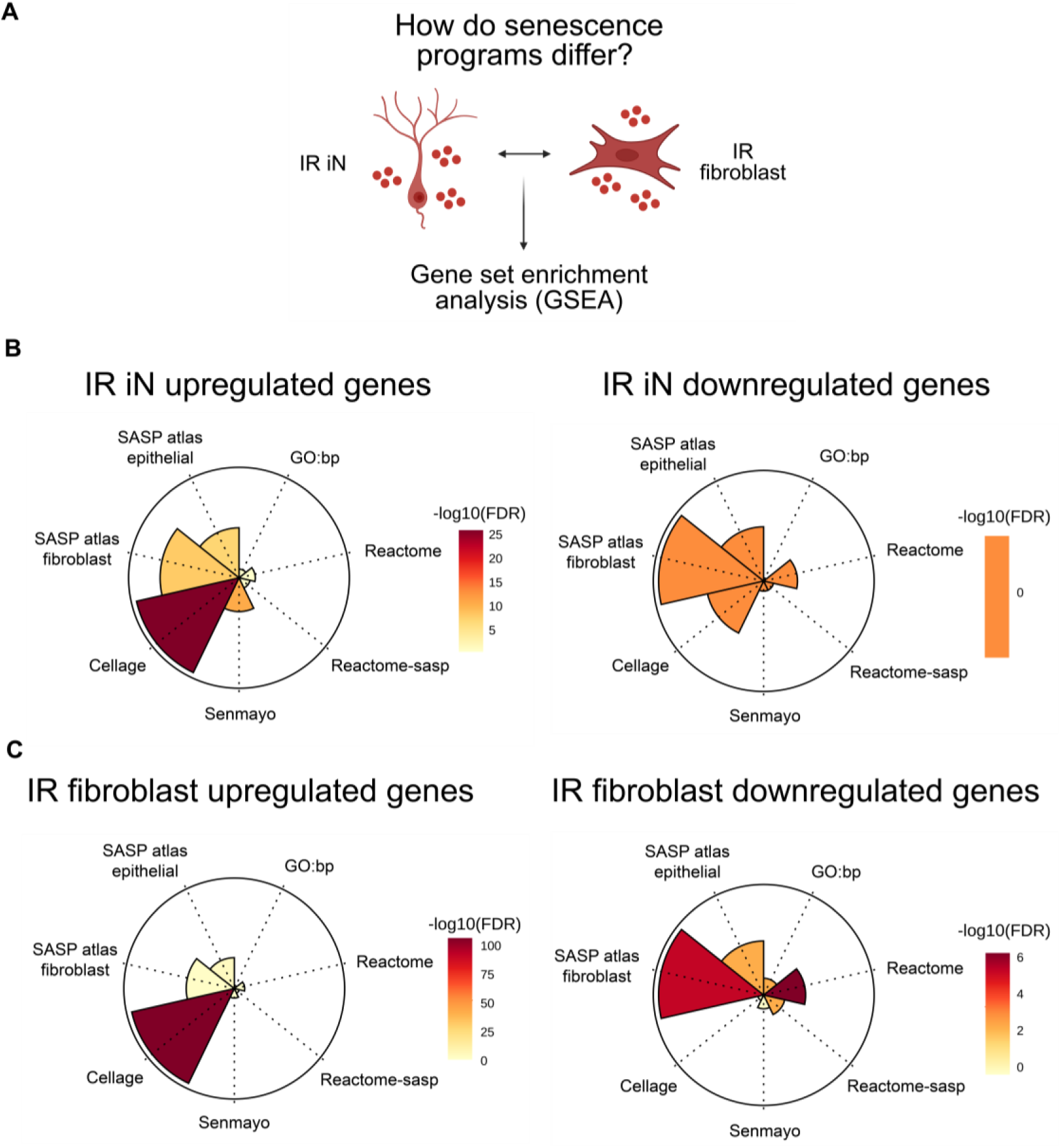
Senescence programs differ between neurons and fibroblasts after DNA damage and diverge from canonical senescence signatures (A) Schematic of using gene set enrichment analysis (GSEA) on IR iN and IR fibroblast RNA-seq data from Fig. 3 (donors: A18, A27, A31, A37, A48, ID97; 1-2 technical replicates per donor in each condition). (B-C) GSEA of senescence-associated datasets. (Left panels) GSEA on upregulated genes. (Right panels) GSEA on downregulated genes.

### Distinct, predicted transcription factor networks underlie DNA damage-induced senescence in neurons and fibroblasts

TFs play key roles in regulating cell fate decisions and may contribute to the differences between IR iN and IR fibroblast senescence-associated cell states. To identify TF networks associated with these responses, we applied a TF activity prediction package curated on a collection of perturbation datasets to infer TF activity from RNA-seq data^36^. This analysis revealed distinct TF programs between the two cell types. IR iNs were predicted to most strongly activate TP53, consistent with activation of DNA damage and stress response pathways (Fig. 5A). In contrast, IR fibroblasts were predicted to strongly repress E2F1, a central regulator of cell-cycle progression, reflecting the changes in proliferative capacity of fibroblasts upon DNA damage, which neurons may not experience (Fig. 5B).

**Figure 5.**
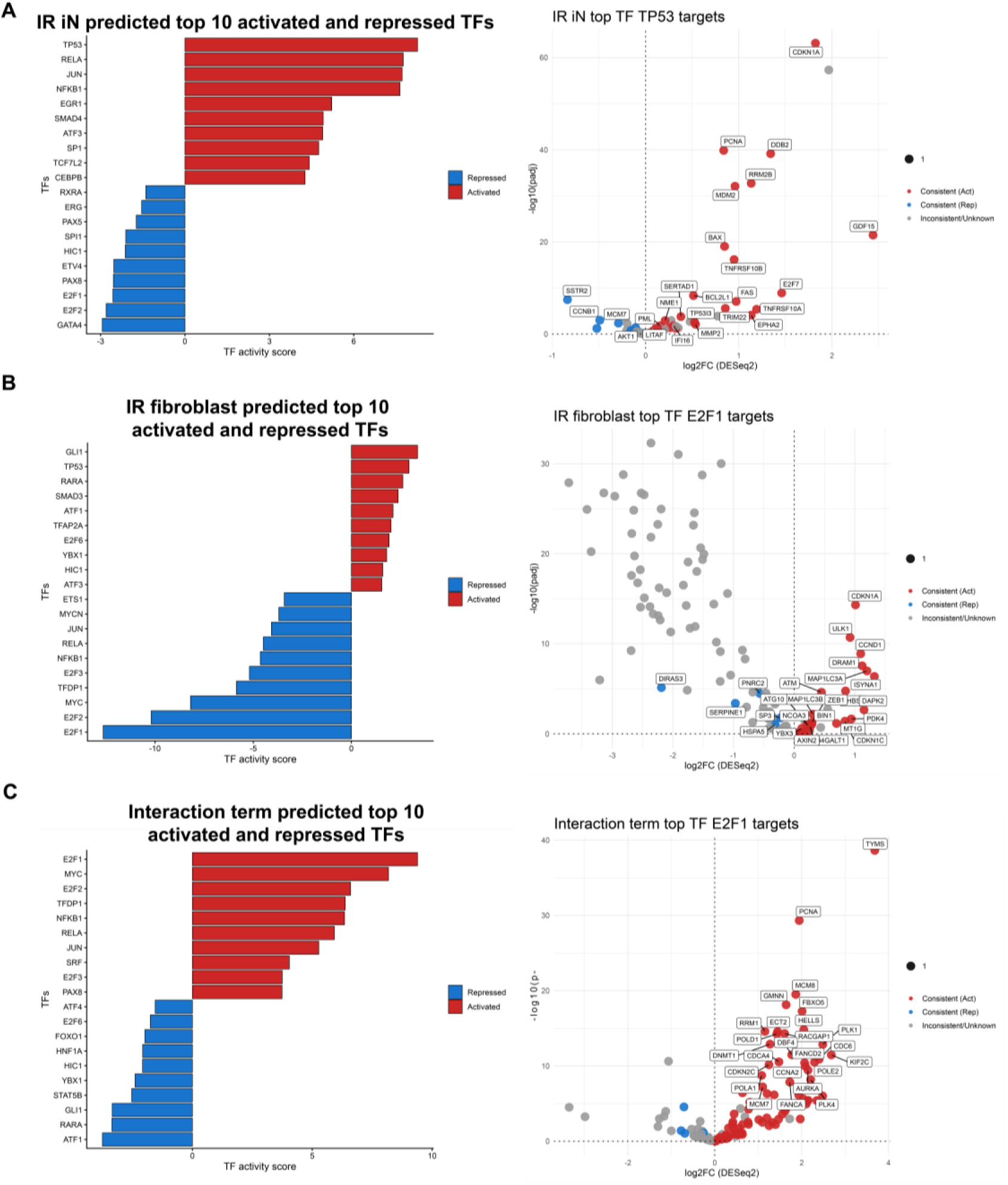
Distinct transcription factor networks underlie DNA damage-induced senescence in neurons and fibroblasts (A-C) Top 10 activated and top 10 repressed transcription factors inferred by decoupleR (ULM) using CollecTRI and DESeq2 shrunken log2FC signatures for RNA-seq data of IR vs CTL in iNs (A), fibroblasts (B), and the interaction term (C) from Fig. 3 (donors: A18, A27, A31, A37, A48, ID97; 1-2 technical replicates per donor in each condition). Bars in the left panels indicate TF activity scores (red, activated; blue, repressed). Right panels display CollecTRI target genes for the top-ranked TF, with points colored by concordance between predicted TF regulation and observed RNA-seq differential expression.

Analysis of the DESeq2 interaction term applied to the TF activity prediction package highlighted cell-cycle regulatory networks as key distinguishing features between neuronal and fibroblast senescent cell states. TFs involved in cell cycle control including E2F1, MYC, and E2F2, were predicted to be strongly repressed in IR fibroblasts compared to IR iNs (Fig. 5C). Of note, NF-κꞴ1 was predicted to be activated in IR iNs but repressed in IR fibroblasts, and was further enriched in the interaction analysis, suggesting that NF-κꞴ1 could be contributing to the differences in SASP gene expression signatures in IR iNs and IR fibroblasts (Fig. 5A-C). This is consistent with the RNA-seq data of IR iNs that showed KEGG enrichment of the NF-κꞴ signaling pathway from Supp. Fig. 1. Together, TF enrichment analysis reveal fundamentally distinct TF networks underlying DNA damage-induced senescence in neurons relative to fibroblasts, uncovering potential targets to modulate these senescence-associated states between cell types.

### Protein level differences in senescence-associated markers further distinguish neuronal and fibroblast responses to DNA damage

mRNA levels do not always correlate with protein abundance, particularly in aging contexts where post-transcriptional regulation can play a significant role^37–39^. Therefore, to determine whether gene expression data were reflected at the protein level, we performed immunocytochemistry (ICC) for established senescence-associated markers in 3 donor-matched iNs and fibroblasts. Cells were stained for p21, p16, LMNB1, HMGB1, γH2AX, and 53BP1, using DAPI staining to measure nuclear area (Fig. 6A). IR iNs showed increased p21 intensity and 53BP1 foci, accompanied by decreased HMGB1 (Fig. 6B, Supp. Fig. 4). In contrast, IR fibroblasts displayed increased p16 intensity and nuclear size, along with decreased p21, LMNB1, and HMGB1 intensity (Fig. 6B-C). LMNB1 and nuclear size did not change significantly in IR-treated iNs, suggesting that neurons may maintain greater nuclear stability after DNA damage compared to fibroblasts (Fig. 6B-C). Taken together, these data further support a model for distinct senescence programs across cell types, where neurons adopt a p21-associated response and fibroblasts exhibit a more canonical p16-associated senescence phenotype.

**Figure 6.**
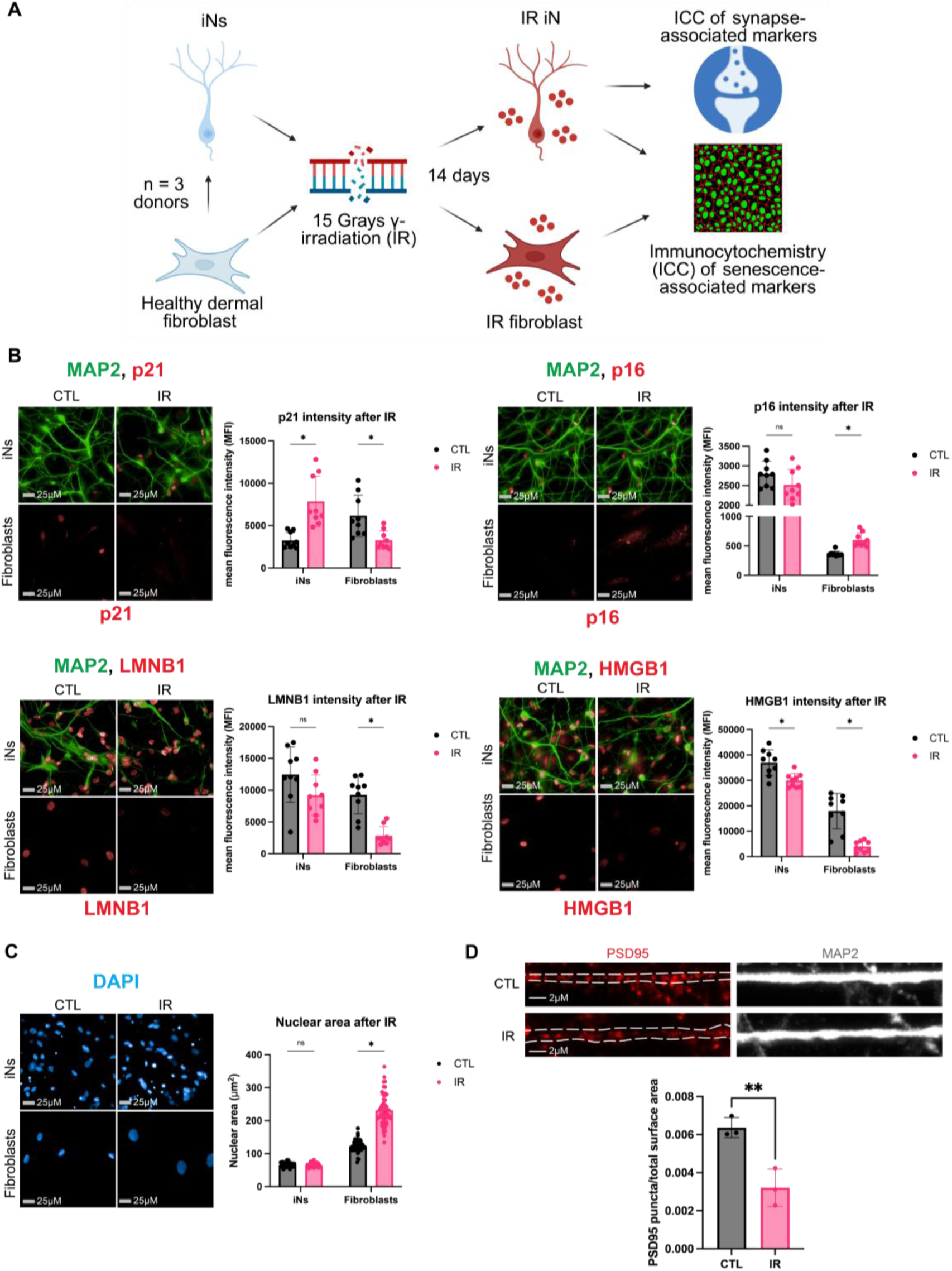
Protein level differences in senescence-associated markers further distinguish neuronal and fibroblast responses to DNA damage (A) Schematic of immunocytochemistry (ICC) performed after IR in healthy dermal fibroblasts and iNs from the same donors (donors: A27, A48, ID98; 3 technical replicates per donor in each condition). 0.2% FBS low serum (LS) treatment was applied to CTL and IR fibroblasts to induce quiescence 72 hours prior to collection. (B) IR iNs and IR fibroblasts were stained for p21 (top left), p16 (top right), LMNB1 (bottom left), and HMGB1 (bottom right). Mean nuclear fluorescence intensity was quantified. (C) Nuclear area of IR iNs and IR fibroblasts quantified by DAPI staining (donors: A27, A48, ID98, 18 technical replicates per donor in each condition). (D) PSD95 staining for IR iNs and IR fibroblasts, imaged by confocal microscopy, and analyzed by Imaris to detect PSD95 spots on MAP2-positive dendrites and normalize to dendrite surface area. Two donors used for PSD95 staining (donors: A27, A48; 3 technical replicates per donor in each condition; donor A48 images and quantification shown in Fig. 6D). Tracks on MAP2 traced in Illustrator. All iNs imaged were quantified with automatic MAP2 tracing to measure p21, p16, LMNB1, HMGB1, and DAPI only in neuronal nuclei or PSD95 only in neuronal dendrites (Supp. Fig. 5). Imaging was performed on the Harmony Operetta CLS system at 63X unless otherwise stated. Nuclear intensity quantification of images performed by the Harmony analysis system and nuclei number after IR provided in Supp. Fig. 4. Statistics were calculated using multiple unpaired t-tests using Welch’s correction with Holm–Šídák adjustment for multiple testing with alpha = 0.05. *p < 0.05, **p < 0.01, ***p < 0.001, ****p < 0.0001, ns: not significant, nd: not detected.

To validate whether synapse-associated dysregulation at the mRNA level translated to the protein level, we stained for PSD95 in the IR iNs and found that PSD95 levels decreased in IR iNs (Fig. 6D), supporting the presence of synaptic dysregulation upon DNA damage in neurons. Although ICC provides semi-quantitative estimates of protein levels, these measurements revealed major differences in the basal levels of senescence-associated proteins before and after IR in iNs and fibroblasts. For example, p16 does not change in IR iNs but is at higher levels than in fibroblasts (Fig. 6B). Additionally, γH2AX and 53BP1 foci were maintained at higher levels in fibroblasts before and after IR than in iNs even as 53BP1 foci increased in IR iNs (Supp. Fig. 4). Differences in the levels of cell cycle inhibitors and DDR proteins at a basal state between neurons and fibroblasts could contribute to which cell cycle regulators are engaged following DNA damage. Overall, our ICC results validate key transcriptomic findings and further support the existence of a distinct DNA damage-induced senescence-associated state in neurons compared with fibroblasts.

### DNA damage induces greater damage lesions and a slower DNA repair-associated response in neurons compared to fibroblasts

Having established fundamental differences in DNA damage-induced senescent states between neurons and fibroblasts, we next investigated whether early DDR responses also diverge between cell types, given their known role in shaping downstream cell fate decisions^20,40–44^.

Consistent with this possibility, we observed that 53BP1 was significantly increased in iNs 14 days after IR but not in IR fibroblasts (Supp. Fig. 4). To investigate further, we carried out a time course ICC experiment to stain iNs and fibroblasts from 2 healthy donors at 0 minutes, 15 minutes, 2 hours, and 24 hours after IR exposure (Fig. 7A). We measured 53BP1 as a proxy for DNA repair by non-homologous end-joining (NHEJ) activity and γH2AX and phosphorylated ATM (pATM) as markers for sites of DNA damage (Fig. 7A)^45–47^. Fibroblasts exhibited a rapid increase in 53BP1 foci approximately 3-fold relative to control at 2 hours, whereas iNs reached a similar increase relative to control only 24 hours after IR (Fig. 7B). In addition, γH2AX foci were significantly elevated 24 hours after IR in iNs compared with fibroblasts that showed only mild increase at this timepoint (Fig. 7C). Further, pATM foci were also increased in iNs 24 hours after IR while fibroblasts did not show pATM foci induction at the same time points measured after IR (Fig. 7D). Total foci number for 53BP1, γH2AX, and pATM, however, was generally higher at each time point after IR in the IR fibroblasts relative to the IR iNs (Fig. 7B-D). These results suggest early differences in DDR between neurons and fibroblasts may lead to faster clearance of DNA damage foci in fibroblasts relative to neurons and be linked to later differences in senescence-associated cell states.

**Figure 7.**
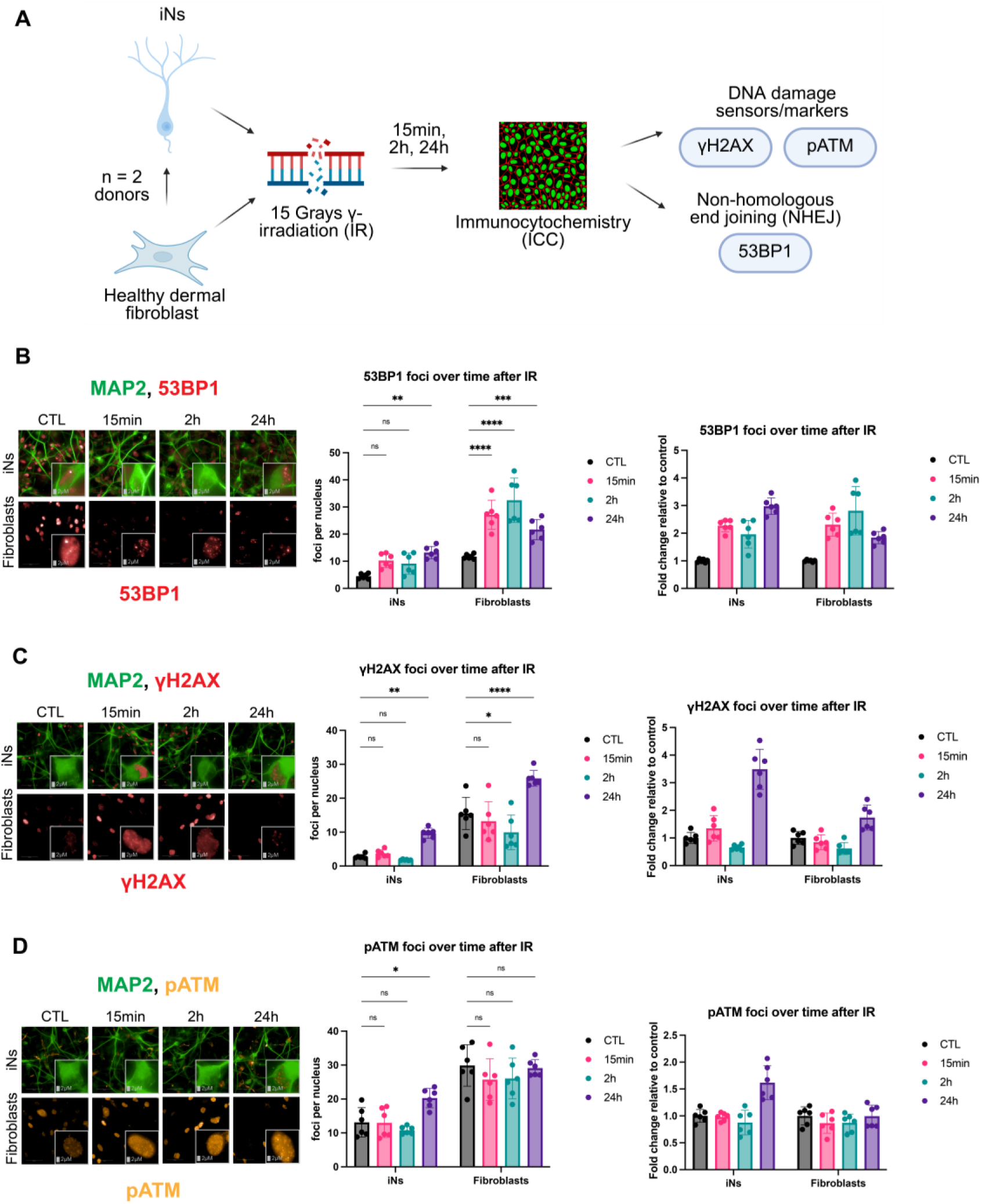
DNA damage induces greater damage lesions and a slower DNA repair-associated response in neurons compared to fibroblasts (A) Schematic of immunocytochemistry (ICC) performed over time after IR in healthy dermal fibroblasts and iNs from the same donors (donors: A48, ID97; 3 technical replicates per donor in each condition). (B-D) iNs and fibroblasts were stained for 53BP1 (B), γH2AX (C), and pATM (D) at 0 min, 15 min, 2h, and 24h after IR with magnified images from the viewfield in the bottom right of each image. Relative fold change (right panels) was calculated from mean foci number per nucleus (middle panels). All iNs imaged were quantified with automatic MAP2 tracing to measure 53BP1, γH2AX, and pATM foci only in neuronal nuclei (Supp. Fig. 5). Imaging was performed on the Harmony Operetta CLS system at 63X. Foci quantification of images performed by the Harmony analysis system. Intensity over time without relative fold change calculations provided in Supp. Fig. 6 along with nuclei number over time after IR. Statistics were calculated by two-way ANOVA (or Mixed Model) and corrected for multiple comparisons using the Dunnett method and reporting multiplicity-adjusted p-value for each comparison with alpha = 0.05. *p < 0.05, **p < 0.01, ***p < 0.001, ****p < 0.0001, ns: not significant, nd: not detected.

## DISCUSSION

Cellular senescence has been increasingly implicated in age-related diseases and more recently in neurodegenerative diseases; however, much of this research has focused on glial cell types in the brain, such as astrocytes and microglia, and the mechanisms driving senescence in neurons remain poorly understood^6–12,48–50^. Here, we demonstrate that DNA damage induces a distinct senescence-like state in neurons that display transcriptional features relevant to AD. Using a direct differentiation system that preserves age-associated transcriptomic and epigenetic signatures, we show that DNA-damaged iNs share transcriptional signatures with published AD iN and post-mortem neuronal datasets, implicating this state in AD pathophysiology. Through WGCNA, we identify both established and novel transcriptional targets of this cell state. Direct comparison with IR-treated fibroblasts from the same donors reveals divergent gene expression profiles, senescence-associated transcriptional states, and predicted transcription factor activity. Critically, these differences extend to the protein level: IR iNs exhibit a p21-associated senescent phenotype while IR fibroblasts adopt a p16-associated phenotype, demonstrating that senescence is not a uniform program across cell types. Finally, temporal profiling of the DDR reveals that neurons exhibit delayed signaling kinetics and sustained accumulation of DNA damage lesions relative to fibroblasts, which may underlie their distinct senescent fate. Together, these findings define a neuron-specific senescent state relevant to AD and reveal fundamental differences in how neurons and fibroblasts perceive and respond to genotoxic stress.

Cellular senescence in neurons is a relatively new concept, in part because senescence has traditionally been defined as a state of irreversible growth arrest, a feature that does not occur in neurons that are largely post-mitotic^1^. However, senescence-associated neuronal phenotypes are increasingly documented in the literature and linked to various neurodegenerative diseases^6–12,48–50^. The first, seminal paper documenting a senescent-like state showed a p21-dependent phenotype driven by the DDR in mice, which aligns with our data showing a p21-associated phenotype in human neurons after genotoxic stress, and to our knowledge, is one of the first studies to document this phenotype in a human context^6^. It remains to be explored which genes are required to drive the IR iN cell state, but we show strong evidence, from the gene expression, WGCNA, and ICC results, that CDKN1A/p21 is one of the driving factors for neuronal senescent-like states after DNA damage. Further mechanistic studies should directly test for a requirement of p21 to validate this hypothesis. In addition, we also performed WGCNA for the IR fibroblasts which showed that vastly different gene modules are driving fibroblast senescent cell states in comparison to IR iNs. Comparing these two datasets and mechanistically dissecting what drives cell-type specific senescence, including differences in SASP enrichment, would also be of high importance to limiting or eliminating the SASP in different cell types across age-related disease contexts.

The vast majority of cells in the body are not cycling at any given time, giving precedence to the study of senescence in non-proliferative cell types to uncover unique and aging-relevant senescent cell states^51–53^. Consistent with this emerging view, our analyses reveal that DNA damage-induced neurons exhibit transcriptional programs associated with senescence while displaying features that differ from canonical fibroblast senescence. One notable distinction between neuronal and fibroblast senescence observed in our study is the differential activation of transcriptional programs. Our differential expression, GSEA, and TF analyses indicate that SASP-associated signatures are enriched in iNs but not fibroblasts under the same senescence-inducing condition. For example, our data predict that iNs robustly activate NF-κꞴ1 after senescence induction, but this response is repressed in fibroblasts. While fibroblasts have been one of the most common cell types to study in the context of cellular senescence and the SASP, our findings suggest that neuronal senescence may involve distinct regulatory programs that could be particularly relevant for neurodegenerative disease contexts such as AD.

It is important to highlight how heterogeneous senescence responses are depending on the cell type, induction method, and microenvironment^1,21,54–57^. While our IR fibroblasts results are different from canonical senescence responses in fibroblasts, much of the foundational research on senescence has been performed using fetal lung fibroblasts as the model cell type^2,22^. Senescence responses are different depending on the starting cell state, so it is likely senescence responses in a younger versus older adult within the same cell type are different as well. In addition, the tissue microenvironment contributes to cells of the same cell type adopting fundamentally different roles depending on the tissue^58^. As a result, even though a fibroblast from the lung and the skin are the same cell type, they will function differently as they inherently need to perform different tissue level roles^59^. Ultimately, our IR fibroblasts results emphasize the need for senescence models to use primary, human cells from the tissue of interest to increase the relevance to human diseases, such as neurodegenerative diseases implicated with senescence.

Although the transcriptional signatures of DNA damage–treated iNs showed strong concordance with previously reported AD neuronal datasets, some differences were observed between our model and previous datasets. The IR iNs displayed a p21-associated cellular state while AD iNs were shown to exhibit a p16-associated cellular state^9^. These findings raise the possibility that neuronal senescence may converge on similar transcriptional programs despite being initiated by distinct upstream triggers. Alternatively, different stages of disease progression or distinct cellular contexts may influence which senescence regulators are engaged. Similarly, while our IR iN transcriptional profile overlapped with neuronal datasets from the entorhinal cortex of AD patients, we observed discordant signatures with the larger ROSMAP dataset. The lack of concordance is mainly from downregulation of synapse-associated genes in the IR iN dataset but upregulation of synapse-associated genes in the ROSMAP AD excitatory neurons. This could be explained by a variety of factors, including differences between a mono-culture and the whole brain environment. It is also possible that synaptic gene expression changes reflect different stages of disease progression. If senescent-like neurons are an early trigger of neurodegeneration, synapse-associated genes could initially decrease due to an increase in senescent cells but then increase as the disease progresses further through compensatory, transcriptional mechanisms. Importantly, proteomics of human AD brains consistently report a decline in synapse-associated proteins with the progression of AD, which aligns with the synaptic decline observed in our IR iN gene expression and protein level data^60^.

The differences in early DDR dynamics observed in neurons and fibroblasts may also contribute to their divergent senescence outcomes. Our time-course analyses reveal delayed recruitment of DNA repair factors and prolonged DNA damage signaling in neurons compared with fibroblasts, suggesting that neurons experience slower or less efficient resolution of DNA lesions following genotoxic stress. Differences in access to DNA repair pathways between proliferative and post-mitotic cells may contribute to these observations. For example, proliferating cells such as fibroblasts can make greater use of homologous recombination during DNA replication, whereas post-mitotic neurons rely more heavily on NHEJ and related repair pathways^61,62^. Thus, future experiments should focus on better characterizing DNA repair access and capacity in neurons and fibroblasts to determine whether DNA repair capacity and DDR signaling influence neuronal senescence. In particular, strategies that enhance DNA repair efficiency in neurons without inducing aberrant cell cycle re-entry may help limit damage accumulation and downstream senescence-associated phenotypes.

Together, our findings demonstrate that DNA damage induces a distinct senescence-like state in neurons characterized by AD-relevant transcriptional signatures, unique regulatory networks, and altered DNA damage responses. These results highlight the importance of studying senescence in post-mitotic cell types and suggest that neuronal senescence may represent a previously underappreciated contributor to neurodegenerative disease pathogenesis.

## METHODS

### Donor information

Primary human dermal fibroblasts were purchased from Lifeline Cell Technology (FC-0024), ATCC (PCS-201-012), or Lonza Bioscience (CC-2511). Primary dermal fibroblasts were also used in this project provided by the University of California Alzheimer’s Disease Research Center (UCI-ADRC) and the Institute for Memory Impairments and Neurological Disorders and isolated at the originating institution through skin punch biopsy. Dermal fibroblasts from UCI were genotyped for APOE status and cognitively diagnosed with AD at the time of biopsy and at a follow-up time period (cognitive follow up performed yearly from the time of enrollment).

Cognitive diagnosis takes into account a comprehensive neurological and physical exam, an extensive medical history with reporting of changes over time, study partner interviews regarding cognitive and behavioral changes, as well as detailed neuropsychological testing (which can last up to 3 hours). The ADRC clinicians then carefully review each case and, if needed, hold a consensus conference to evaluate all the information described above and determine the diagnosis for that visit. If all areas are within the normal range for an individual’s age, education, language, and related factors, the participant may be classified as cognitively unimpaired. A complete list of details for each donor fibroblast can be found in the “Supplementary Table 1 - Data Donor Information.”

### Cell culture

Dermal fibroblasts were cultured in DMEM (Corning, 01-017-CV) supplemented with 15% tetracycline-free FBS (R&D Systems, 631101), 1X Antibiotic-Antimycotic (Life Technologies, 15240062), and 1X MEM Non-Essential Amino Acids Solution (Life technologies, 11140050) at 37°C, 3% O2, 5% CO2. All fibroblasts were cultured with the same lot of FBS (Lot number 241171501). 3% O2 was used for culturing conditions to match physiological oxygen levels in the skin and brain^63–65^. Primary dermal fibroblasts were cultured in 12-well plates (Genesee Scientific, 25-100) for sequencing experiments or 96-well microplates (VWR, MSPP-P9615HN) for microscopy imaging, with media changes every 2–3 days. All cultured cells were monitored and tested regularly for mycoplasma (Lonza, LT07-318). All cultured cells were counted at each split and population doublings (PD) were monitored over the course of the culture. PDs were kept below 30 for all experiments with no noticeable effects on proliferation before or after PD 30 for all primary cells. PDs were matched between CTL and IR samples when possible. Healthy donors were between ages 18-73. Donor samples were not sex-matched.

### Patient-derived directly-induced neurons (iNs)

Dermal fibroblasts were transduced by lentivirus expressing inducible human NGN2 and ASCL1 (plasmid pLVX-UbC-rtTA-Ngn2:2A:Ascl1 (Addgene, 127289)) for 24 hours at half the normal culture media volume in normal growth media with 4µg/mL polybrene at a multiplicity of infection (MOI) of 20 (Sigma, TR-1003-G)^66^. A relatively high MOI was used, as previously described, for high, inducible expression of human NGN2 and ASCL1 to facilitate fibroblast to neuron conversion without noticeable effects on proliferation^66^. Media was refreshed after transduction and selection started 72 hours later with 1µg/mL puromycin. After transduction, dermal fibroblasts were always cultured with selection reagent until conversion without noticeable effects on proliferation. Conversion of fibroblast to neuron started after seeding the transduced fibroblasts at 3X confluency and switching to conversion media for 3 weeks with media changes every 2-3 days. After 3 weeks, iNs were switched to maturation media for a maximum of 2 weeks. iNs were not sorted at the 3 week conversion timepoint or cocultured with astrocytes, unless stated otherwise. Sorting was previously performed to enrich neuronal signatures; however, validation of the bulk RNA-seq iN cultures without sorting has been performed through direct comparison to pure fibroblast cultures and protein staining specifically in neurons to validate gene expression results^12^. Details on the conversion process and conversion and maturation media formulations have been described previously^12,67^.

### Senescence induction and drug testing

Senescence was induced with IR. For IR-induced senescence, non-transduced fibroblasts were irradiated at 25-50% confluency with 15 Grays (Gy) and iNs were irradiated at the 3 week conversion time point. Both cell types had medium change performed immediately after IR. Cells were considered senescent after 14 days post IR, during which medium was regularly changed (every 2–3 days). Fibroblasts were cultured in full serum throughout the senescence time period unless stated otherwise. 0.2% FBS low serum (LS) treatment was applied to induce quiescence in Fig. 6 ICC results for both CTL and IR fibroblast conditions as previously described^22^. LS treatment was applied 72 hours prior to collection with a wash in LS 72 hours prior to collection and replacement with LS media, followed by a final refresh 24 hours prior to collection.

### Bulk RNA-seq processing

Bulk RNA-seq libraries were generated from total RNA isolated from samples collected at 0h and 336h (14 days) post IR. Mock IR non-transduced fibroblasts were collected at 0h post IR while mock IR iNs were collected at 336h to match differentiation timelines in the CTL iNs, which is not present in the non-transduced fibroblasts. The cells were lysed in the lysis buffer plus 2-Mercaptoethanol (Sigma, M3148-25ML) according to manufacturer’s instructions (VWR, BIO-52073). RNA was isolated according to the protocol and sent out to Novogene for sample QC, library preparation, and sequencing. Sequencing was performed at 20M paired-end reads per sample on the NovaSeq X Plus Series (PE150) (Illumina). 1-2 technical replicates per donor from CTL and IR conditions were sequenced. FASTQ files provided by the vendor were used as the starting point for computational analyses.

Raw reads were aligned locally using the nf-core/rnaseq salmon pipeline in Windows Subsystem for Linux (WSL), performing pseudo-alignment against a standard reference transcriptome for Homo sapiens (GRCh38 genome)^68,69^. Transcript-level count estimates (estimated counts and TPMs) per sample were then summarized to gene-level count matrices using the corresponding transcript-to-gene annotation in R. The gene-level estimated count matrices along with sample metadata were used for bulk RNA-seq dataset generation and analyses. Sample metadata can be found in this table: “Supplementary Table 2 - Combined Metadata.”

### Differential expression analysis

DESeq2 formula used is provided here for fibroblasts or iNs: ∼ batch + condition^70^. DESeq2 formula used for the interaction term between fibroblasts and iNs is provided here: ∼ batch + cell.type * condition. Principal component analysis (PCA) provided in Supp. Fig. 7A showing rationale to batch correct iN samples. Top 10 up and down DEG heatmap is provided in Supp. Fig. 7B to show that top dysregulated genes are aligned across samples after batch correction. Supp. Fig. 7C gives the rationale from the PCA to include cell.type and condition when computing the DESeq2 interaction term. DEG cutoffs were defined by |log2FC| > 0.5 and adjusted p-value < 0.05. IR iN versus IR fibroblast upregulated and downregulated DEG venn diagrams provided in Supp. Fig. 3. For volcano plots, the top 10 DEGs by significance are labeled and the x-axis is limited to a log2FC range of −5 to 5. GO and KEGG enrichment performed as previously described^34,70–72^. KEGG enrichment provided in Supp. Fig. 1. The RRHO2 package was used for RRHO comparisons^24^. TF enrichment was performed using the Collection of Transcription Regulation Interactions meta-resource that was recently sign-curated for activation or repression of each TF in the database^36^. DEG lists can be found in “iN_IR_vs_CTL_ALL.csv,” “fibro_IR_vs_CTL_ALL.csv,” and “interaction_iN_minus_fibro_ALL.csv.”

### Public databases used for RRHO comparisons

IR iNs were compared by RRHO to AD iNs, scRNA-seq data from the entorhinal cortex of AD patients, and scRNA-seq data from the prefrontal cortex of AD patients (ROSMAP). The AD iN and entorhinal cortex datasets were downloaded from publicly available sources and the ROSMAP dataset was downloaded from the AD Knowledge Portal. The results published here are in whole or in part based on data obtained from The AD Knowledge Portal (https://doi.org/10.7303/9618239).

Prior to running DESeq2 on the publicly available AD datasets, scRNA-seq data was pseudobulked based on neuronal cell type and the ROSMAP scRNA-seq data was specifically pseudobulked on excitatory neurons prior to RRHO comparison. The AD dataset DESeq2 formula was based on the condition of AD and for the ROSMAP dataset, the DESeq2 formula was based on the Braak, Cerad, or Cogdx scores.

### Immunostaining

Cells were fixed with 4% paraformaldehyde for 15 min, and then washed three times with PBS followed by permeabilization and blocking with 10% normal goat serum (Life Technologies, 50062Z) in 0.1% Triton X-100 (Sigma-Aldrich, X100RS-5G) for 1h. Cells were then incubated with primary antibodies at 4°C overnight (γH2AX, abcam, ab81299, 1:250; 1:1000; pATM, Life Technologies, MA1-2020, 1:250; p21, abcam, ab109520, 1:100; p16, abcam, ab270058, 1:50; LMNB1, abcam, ab229025, 1:1000; HMGB1, abcam, ab79823, 1:250; 53BP1, abcam, ab175933, 1:200, PSD95, Life Technologies, MA1-046, 1:300; MAP2, Novus Biologicals, NB300-213, 1:400). The next day, after washing three times with PBS for 5 min each wash, secondary antibodies were applied for 1 hour. Cells were then washed once with PBS, counterstained with DAPI (Sigma, D9542-1MG) at 0.5 μg/mL in MQ water for 15 minutes, and washed once with MQ water. Finally, the cell solution was replaced with PBS before imaging. All steps were performed at room temperature except for primary antibody incubation overnight.

### Image acquisition and analysis

Cells were plated on glass-based 96-well plates. Images were taken on Harmony Operetta CLS type HH1600. Images were taken at 63X with water immersion, NA 1.15, and two peak autofocus. Confocal imaging was not used unless stated otherwise. Camera ROI was set to 2160x2160. Binning was set to 2. Images were captured using the following channels: DAPI, Alexa 488, Alexa 555, and Alexa 647. For each ICC experiment, 3 technical replicates per donor were imaged spread across 3 wells and 3-6 tiles per well were captured. 11 stacks per tile were imaged at a 0.4μm distance between stacks. Fig. 6 results imaged and analyzed an average of 47-57 nuclei for the iNs and 16-30 nuclei for the fibroblasts across conditions. Fig. 7 results imaged and analyzed an average of 136-150 nuclei for the iNs and 128-139 nuclei for the fibroblasts across conditions.

Images analyzed with the pipeline as described here on the Harmony analysis software. Flatfield correction set to basic, Stack processing set to “maximum projection”. Nuclei segmented using DAPI channel, Harmony method A with common threshold set to 0.05 and minimum area 15μm^2^. Background subtraction for each channel performed with formula A-quantile(A, 0.10).quantile. MAP2 neurons were defined using “Find Cells” with channel set to MAP2 and using Method M, with diameter set to 10μm, splitting sensitivity=0.00 and common threshold=0.14. Neuronal nuclei were defined as nuclei within MAP2 positive cells using “Find nuclei” with DAPI channel, with region of interest set to MAP2 neurons, using method A with common threshold 0.05 and area greater than 15μm^2^. Neuronal cytoplasm was defined using “Find cytoplasm” with the MAP2 channel, based on neuronal nuclei, using Method B with common threshold=0.65, individual threshold=0.30. Cell and nuclear areas were calculated using “Calculate Morphology Properties” on the indicated population with Method set to “Standard”. Neurites were defined using “Find Neurites” based on MAP2 positivity from the “Neuronal Nuclei” population with the region set to “Nucleus” and with method “CSIRO Neurite Analysis 2” with smoothing width = 3px, linear window = 20px, contrast >5, diameter >= 7px, gap closure distance <= 7px, gap closure quality =0, debarb length <= 15px, body thickening = 5px, tree length <= 20px. Intensities for neuronal stains were calculated using “Calculate intensity properties” for the population “Neuronal nuclei”, region “nucleus”, using the method “Standard” with output “mean”. Neuronal intensities were calculated using “calculate intensity properties” for the population "neuronal nuclei” region “cell” with the method “standard” and output “mean”. Spots/foci/puncta were defined using “find spots” with ROI set to Neuronal Nuclei, region either set to “nucleus”, “neurite”, or “cytoplasm”, using method “C” with radius <= 1.18 um, contrast > 0.10, and uncorrected spot to region intensity > 1.0. Colocalization of puncta was defined using “find spots” with the ROI for one of the stains of interest set to previously defined puncta of the other marker of interest. Spots per area were calculated by dividing the quantity of puncta per well by the sum of the area of neuronal cytoplasm (or neurite length) per well. Supp. Fig. 5 shows example images of the neuronal segmentation pipeline to calculate intensities and foci only in neuronal nuclei.

### Weighted gene co-expression network analysis (WGCNA)

WGCNA was run using pyWGCNA^73^. The topological overlap matrix similarity dendrogram was made using the R WGCNA package and the gene matrices were made using the BioMart R package^27^. The gene expression matrix used for pyWGCNA was batch-corrected and ran through a variance stabilizing transformation which was then filtered for top 75% most varying genes. This was run for both the IR iN and IR fibroblast datasets separately (IR fibroblast WGCNA results provided in Supp. Fig. 8). Resulting gene modules were tested for gene set enrichment analysis (GSEA) against GO Terms (Gene Ontology), as previously described, and cross-referenced with differentially expressed genes (DEG)^34^. Our iN sample number gave us confidence that the WGCNA would reach sufficient power for meaningful results, as Langfelder and Horvath recommend using at 15 samples and our total sample number was 18 and 1-2 biological replicates per sample^27^. It should also be noted that meaningful results can still be discovered with less than 15 samples, which is where our fibroblast sample number fell below, and emphasizes the importance of validating WGCNA module genes regardless of sample size^74^. Supp. Fig. 3 shows module genes being compared to DEGs from IR iN and IR fibroblasts genes and overlap lists provided in “_overlap.csv” supplementary tables. WGCNA module names are randomly generated by the pyWGCNA package and do not have a biological meaning.

### Gene set enrichment analysis (GSEA)

Gene enrichment was run using the GeneOverlap package (doi:10.18129/B9.bioc.GeneOverlap) on the significantly up and down DEGs and GSEA was run from the clusterProfiler package on the full list DEGs. Enrichment and GSEA were completed using gene sets for hallmarks of aging and senescence-specific genes from CellAge, SenMayo, SASP Atlas (fibroblast and epithelial), REACTOME, and GO^22,29–35^.

### Confocal synapse imaging and quantification

For imaging of synapses, the Zeiss LSM 980 laser scanning confocal microscope on an inverted Axio Observer 7 SP microscope was used to image iNs on glass-based 96-well plates. Images were taken at 63X with a scanning speed of 6 plus 3-5 stacks at 0.5μm intervals. 1 view field across 3 wells for 2 donors (A27, A48) was imaged. Image quantification was performed in Imaris by segmenting at least 5 secondary synapses per condition, view field, and donor based on MAP2 and masking the surface to then find spots on the MAP2 surface^75^. The total number of spots were then divided by the total surface area of surface regions of interest. Images were processed in Fiji and pseudocolored for clarity^76^. The dotted tracks along the synapse and scale bar were made and adjusted, respectively, in Illustrator.

### Code availability

Code will be made available on github upon acceptance of manuscript.

## DECLARATIONS

### Competing interests policy

No competing interests to report.

### Data availability

Raw sequencing data (FASTQ files) and processed bulk RNA-seq generated in this study will be deposited in the Gene Expression Omnibus (GEO); accession numbers will be provided upon acceptance of the manuscript. Combined metadata will also be made available upon final manuscript submission. All other data used in this study were obtained from previously published, publicly accessible resources, with source identifiers specified in the manuscript.

## Supporting information

Supplementary Table 2 - Combined Metadata

Supplementary Table 1 - Donor Information

DESeq2_iN_tp336_normal_IR_vs_CTL_allgenes_degs__coral__overlap

DESeq2_iN_tp336_normal_IR_vs_CTL_allgenes_degs__gainsboro__overlap

DESeq2_iN_tp336_normal_IR_vs_CTL_allgenes_degs__salmon__overlap

fibro_IR_vs_CTL_ALL

iN_IR_vs_CTL_ALL

interaction_iN_minus_fibro_ALL

## Acknowledgements

We would like to acknowledge Dr. Judith Campisi and the legacy she built in the field of senescence. We would like to thank the Buck Institute for letting the lab continue until we finish publishing papers started by Judy. We would also like to acknowledge UCI for providing donor fibroblasts to the Buck and the Gage lab for guidance with the iN differentiation. All schematics/diagrams were made using BioRender. The UCI-ADRC is funded by NIH/NIA Grant P30 AG066519. This work was partially supported by P01AG066591 and Training Grant - NIA T32 AG052374. Study data were provided by the Rush Alzheimer’s Disease Center, Rush University Medical Center, Chicago. Data collection was supported through funding by NIA grants P30AG10161 (ROS), R01AG15819 (ROSMAP; genomics and RNAseq), R01AG17917 (MAP), R01AG30146, R01AG36042 (5hC methylation, ATACseq), RC2AG036547 (H3K9Ac), R01AG36836 (RNAseq), R01AG48015 (monocyte RNAseq) RF1AG57473 (single nucleus RNAseq), U01AG32984 (genomic and whole exome sequencing), U01AG46152 (ROSMAP AMP-AD, targeted proteomics), U01AG46161(TMT proteomics), U01AG61356 (whole genome sequencing, targeted proteomics, ROSMAP AMP-AD), P30AG072975, the Illinois Department of Public Health (ROSMAP), and the Translational Genomics Research Institute (genomic). Additional phenotypic data can be requested at www.radc.rush.edu. This work was partially supported by the National Institute on Aging (NIA) 4 RF1 AG056306-07; California Institute for Regenerative Medicine (CIRM) grant INFR6.2-15440 (FHG).

## Research funding

Jun-Wei B. Hughes, National Institute on Aging, T32AG052374

Judith Campisi and Lisa Ellerby (MPI), National Institute on Aging, P01AG066591

Ryo Higuchi-Sanabria, National Institute on Aging, R01AG079806

Fred H. Gage, The Salk Institute for Biological Studies, 4 RF1 AG056306-07

## AUTHOR CONTRIBUTIONS

JWB contributed to all parts of the experiments, analysis, and manuscript. AS and DC performed cell culture, ICC, imaging, analysis, and edited the manuscript. FS and KS helped with bioinformatics analyses including RRHO, GSEA, and WGCNA. RB helped with cell culture and RNA collection. TLMM helped with synapse imaging and quantification. IB provided support on the CLS HTS. HD helped with RNA collection. TAUH provided the base WGCNA code. SM and KAW cultured stocks of donor fibroblasts from UCI at the Buck. HD and MBJ cultured and provided stocks of donor fibroblasts from UCI. JH provided initial starting materials to setup the iN platform at the Buck. RHS edited the manuscript. FHG oversaw the collaboration with the Salk. DF runs the bioinformatics core at the Buck where KS and FS performed analyses. LME provided support on all aspects of the projects including setting up receiving donor fibroblasts from UCI. PYD oversaw and directly supervised all aspects of the project. JC led the project until her passing in January 2024. All authors reviewed the manuscript.

