## Supplementary Data for "DNA damage drives a unique, Alzheimer’s disease-relevant senescent state in neurons"

Supplemental Figure 1. KEGG enrichment of iNs and fibroblasts after IR.

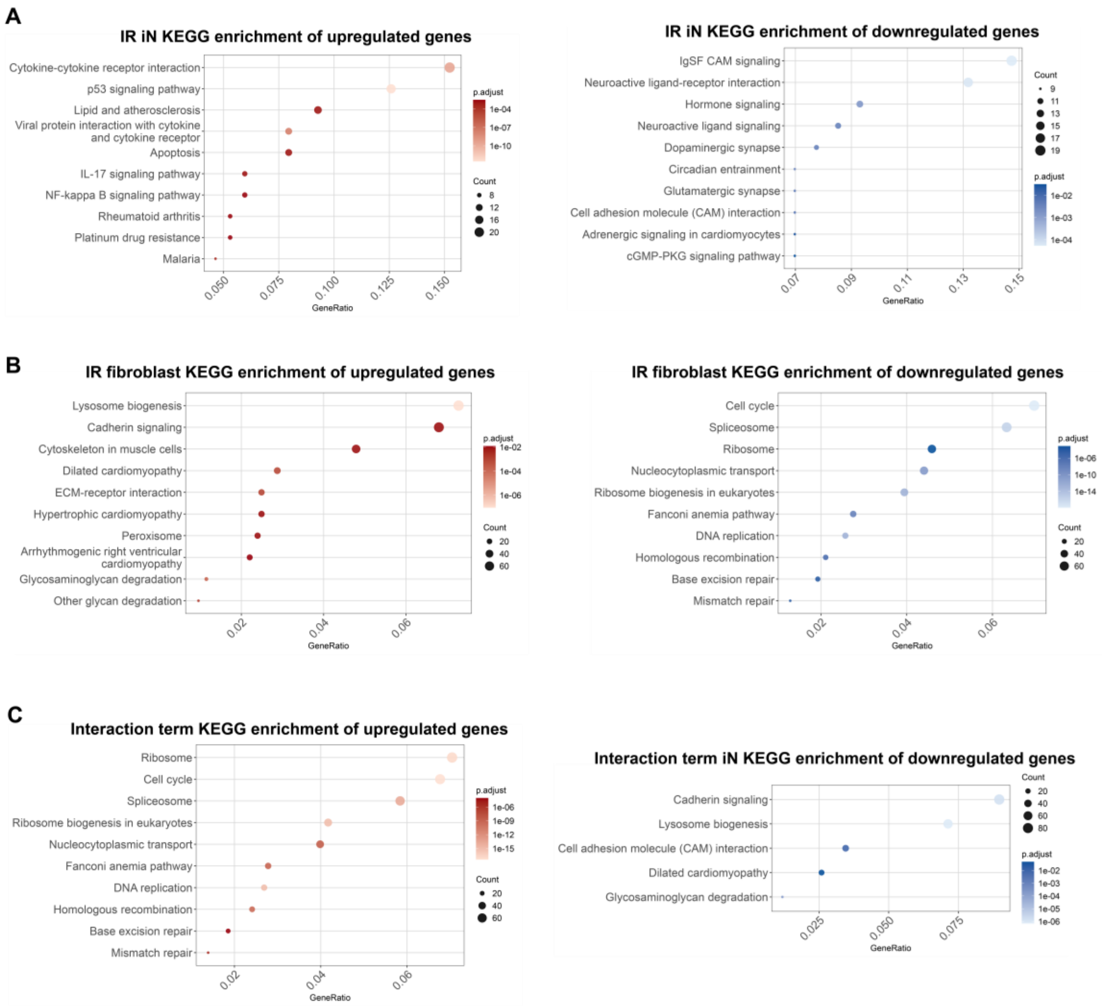


(A-C) Top 10 enriched KEGG terms of upregulated (left panels) and downregulated (right panels) DEGs from IR iNs (A), IR fibroblasts (B), and the interaction term (C).

Supplemental Figure 2. DNA-damaged neurons and ROSMAP Alzheimer's disease neurons display discordant gene expression patterns.

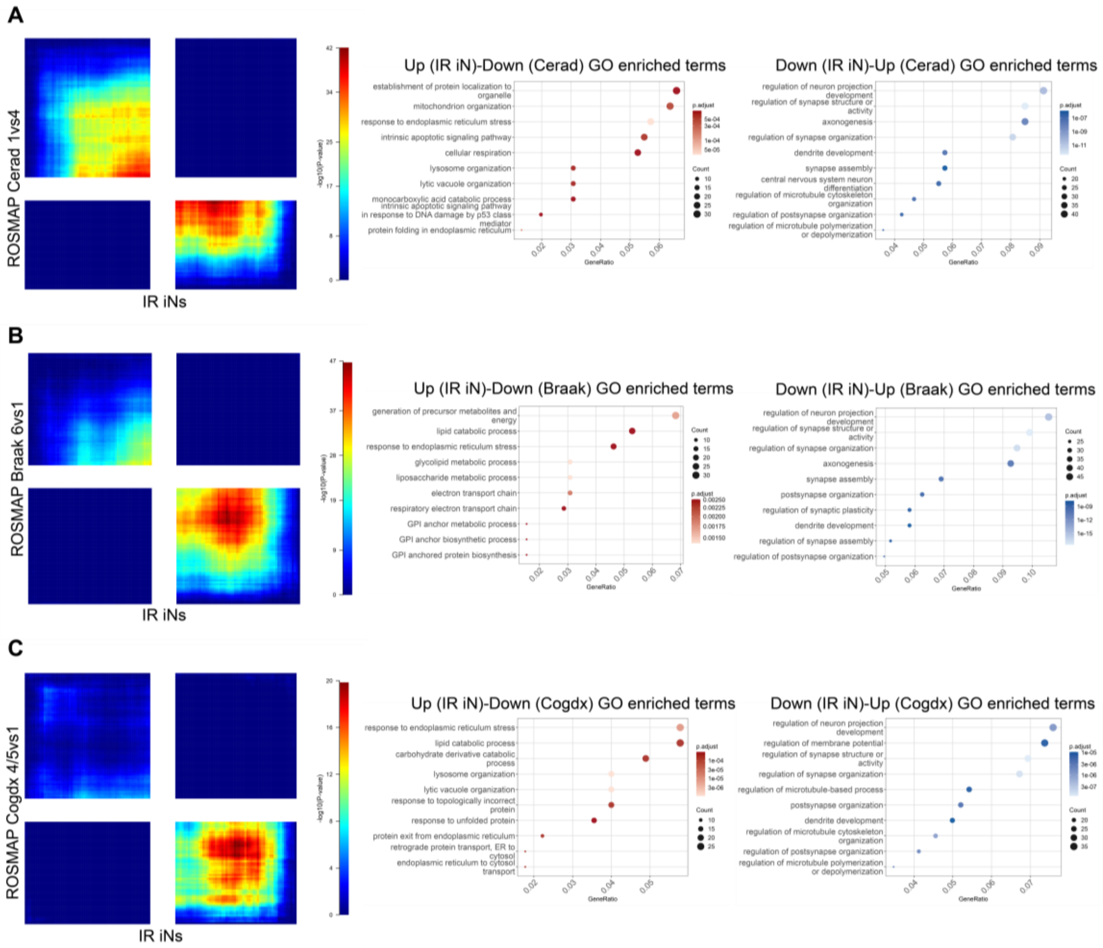


(A-C) Left panels show RRHO plots of IR iN DEGs compared to pseudobulked excitatory AD neurons from ROSMAP broken down into Cerad (A), Braak (B), and Cogdx (C) comparisons. Middle and right panels show top 10 GO enrichment terms provided for discordant quadrants in each comparative study. The bottom right quadrant of the RRHO indicates discordant genes that are upregulated genes in the IR iN dataset but downregulated in the ROSMAP dataset for that condition. The top left quadrant of the RRHO indicates discordant genes that are downregulated in the IR iN dataset but upregulated in the ROSMAP dataset for that condition.

Supplemental Figure 3. Overlapping module genes and differentially expressed genes in both IR iN and IR fibroblasts datasets.

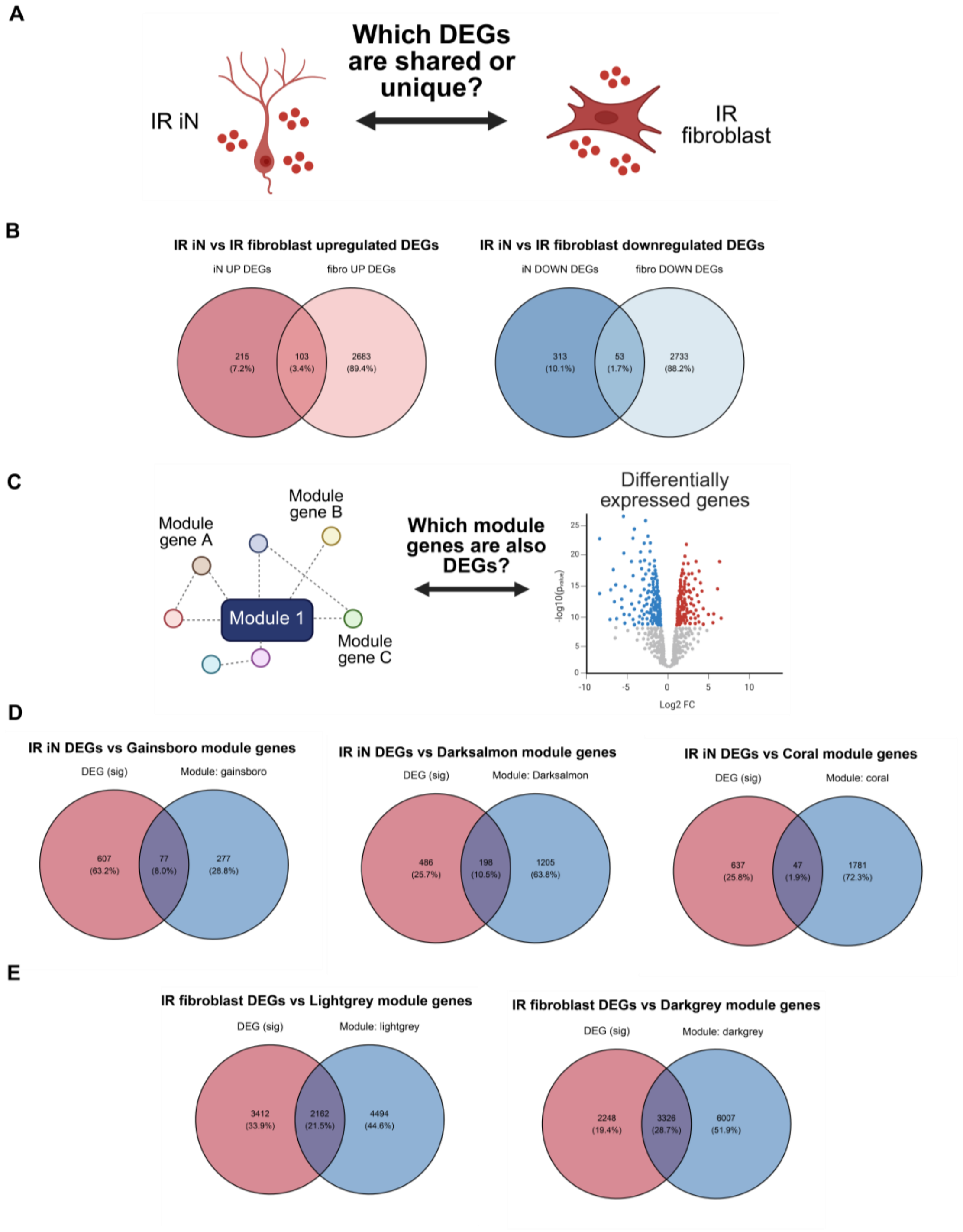

(A) Schematic of comparing IR iN DEGs to IR fibroblast DEGs.

(B) Venn diagrams comparing IR iN versus IR fibroblast DEGs. Upregulated DEG comparison (left) and downregulated DEG comparison (right).

(C) Schematic of comparing module genes to DEGs.

(D-E) Venn diagrams of overlap between module genes and significant DEGs and significant DEGs. IR iN DEGs compared to the Gainsboro (D, left), Salmon (D, middle), and Coral (D, right) modules from the IR iN WGCNA in Fig. 2. IR fibroblast DEGs compared to the Lightgrey (E, left) and Darkgrey (E, right) modules from the IR fibroblast WGCNA in Supp. Fig. 8.

Supplemental Figure 4. Protein level differences in damage-associated markers between neurons and fibroblasts induced to senesce by DNA damage.

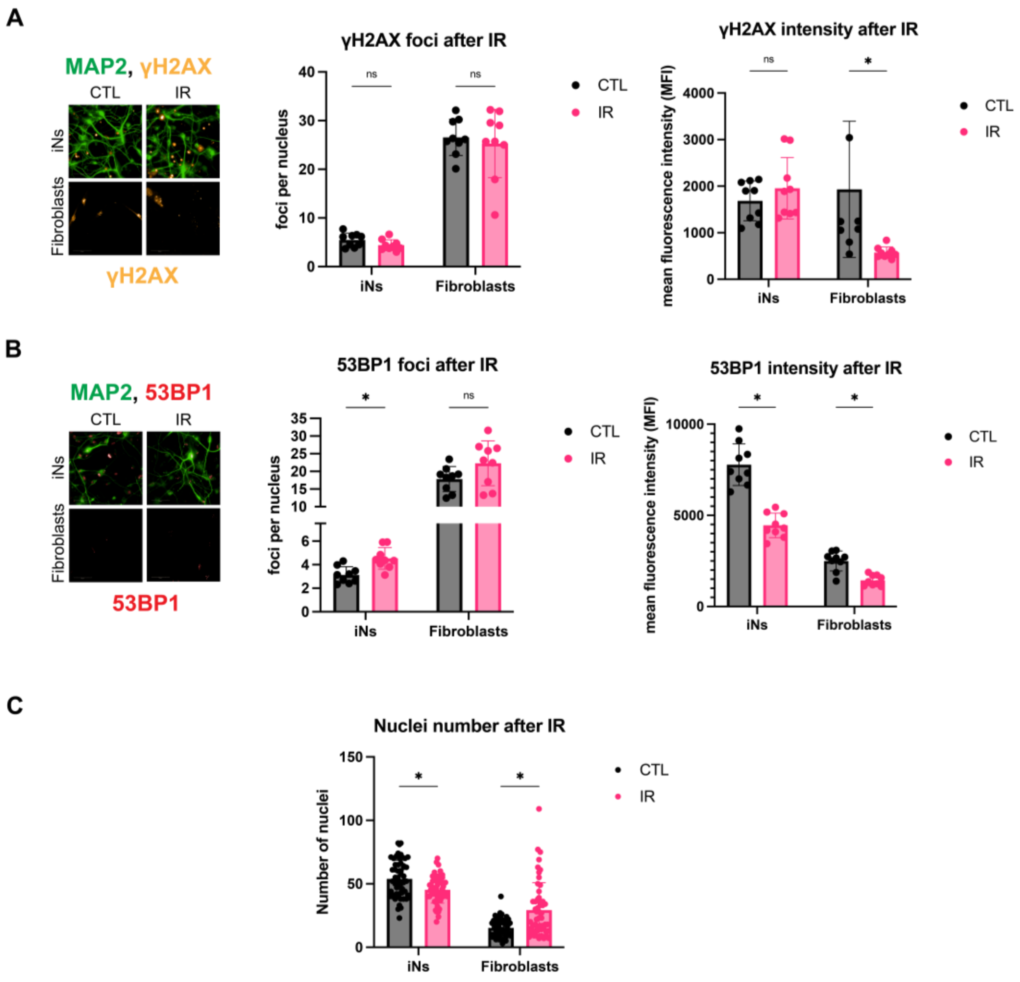

(A-B) Healthy dermal fibroblasts and iNs from the same donors (donors: A27, A48, ID98; 3 technical replicates per donor in each condition) exposed to IR and stained for γH2AX and 53BP1.

(C) Nuclei number quantified by DAPI staining in IR iNs and IR fibroblasts (donors: A27, A48, ID98; 18 technical replicates per donor in each condition). All iNs imaged were quantified with automatic MAP2 tracing to measure γH2AX and 53BP1 intensity and foci only in neuronal nuclei (Supp. Fig. 5). All images were taken on the Harmony Operetta CLS system at 63X. Intensity and foci quantification of images was performed by the Harmony analysis system. 0.2% FBS low serum (LS) treatment was applied to CTL and IR fibroblasts to induce quiescence 72 hours prior to collection. Statistics were calculated using multiple unpaired t-tests using Welch's correction with Holm–Šídák adjustment for multiple testing with alpha = 0.05. *$p < 0.05$, **$p < 0.01$, ***$p < 0.001$, ****$p < 0.0001$, ns: not significant, nd: not detected.

Supplemental Figure 5. High-throughput imaging and analysis pipeline to quantify neuronal nuclei immunocytochemistry staining.

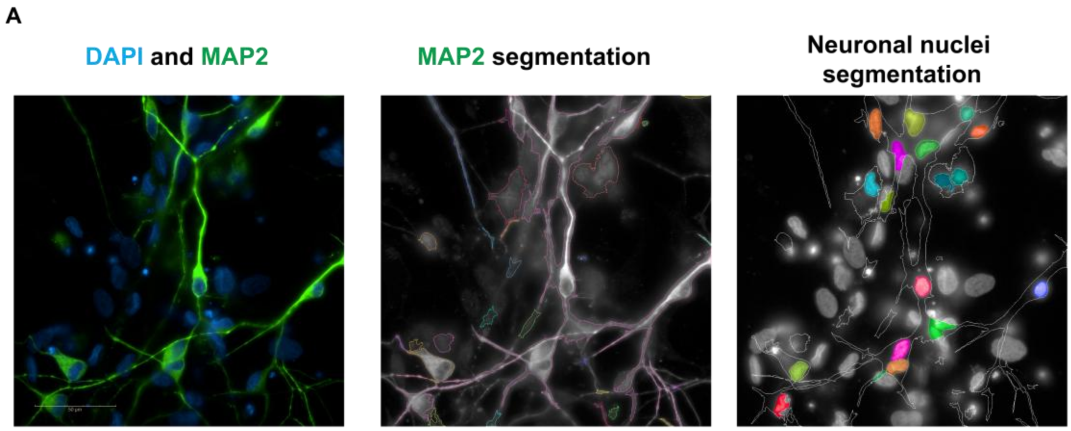


(A) Harmony automated analysis system used to trace neuronal dendrites by MAP2 (left panel) using neuronal segmentation pipelines (middle panel) to determine staining in specifically neuronal nuclei (right panel). The full Harmony pipeline can be found in the “Image acquisition and analysis” section of Methods.

Supplemental Figure 6. DNA damage signaling and DNA repair-associated response in neurons and fibroblasts over time after IR.

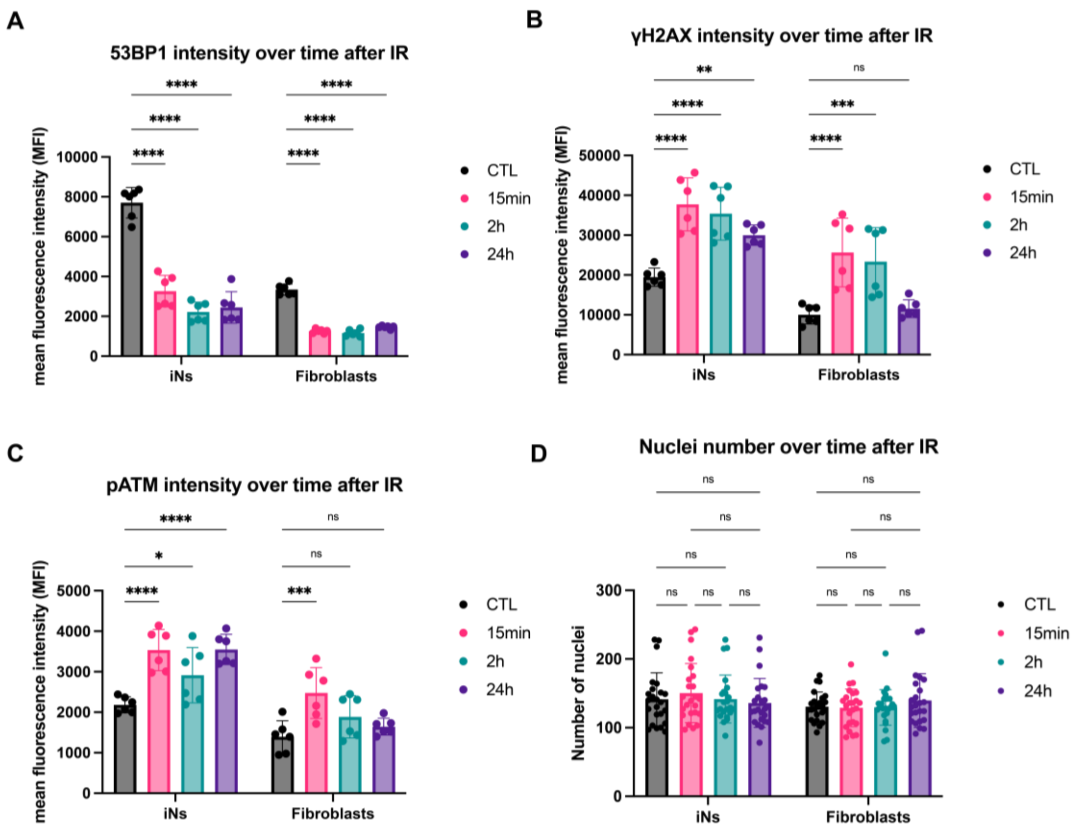

(A-C) ICC performed over time after IR in healthy dermal fibroblasts and iNs from the same donors (donors: A48, ID97; 3 technical replicates per donor in each condition). iNs and fibroblasts were stained for 53BP1 (A), γH2AX (B), and pATM (C) at 0min, 15min, 2h, and 24h after IR.

(D) Nuclei number quantified by DAPI staining in IR iNs and IR fibroblasts (donors: A48, ID97; 12 technical replicates per donor in each condition). All iNs imaged were quantified with automatic MAP2 tracing to measure 53BP1, γH2AX, and pATM intensity and foci only in neuronal nuclei (Supp. Fig. 5). All images were taken on the Harmony Operetta CLS system at 63X. Intensity and foci quantification of images was performed by the Harmony analysis system. Statistics were calculated by two-way ANOVA (or Mixed Model) and corrected for multiple comparisons using the Dunnett method and reporting multiplicity-adjusted p-value for each comparison with alpha = 0.05. *$p < 0.05$, **$p < 0.01$, ***$p < 0.001$, ****$p < 0.0001$, ns: not significant, nd: not detected.

Supplemental Figure 7. Validation of DESeq2 formula for subsequent differential expression analysis.

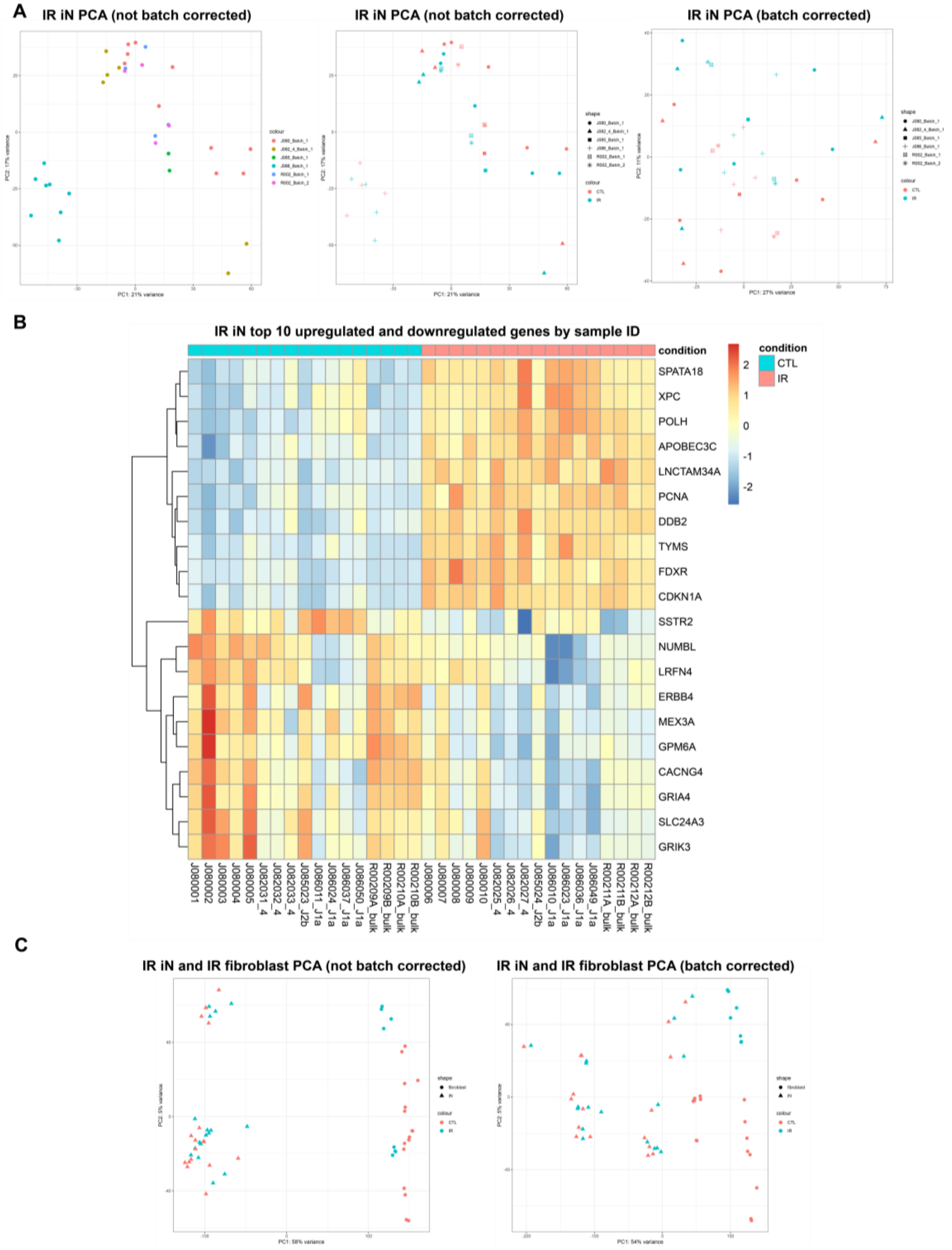

(A) Principal component analysis (PCA) of iNs by batch (left), batch and condition (middle), and batch-corrected condition (right).

(B) Heatmap of top 10 upregulated and downregulated genes in CTL and IR iNs ranked by adjusted p-value.

(C) PCA of interaction term with both iNs and fibroblasts showing condition and cell type versus the PCA of the batch-corrected interaction term.

Supplemental Figure 8. Weighted gene correlation network analysis of fibroblasts after IR shows highly connected modules related to the cell cycle, DNA repair, and organelle function.

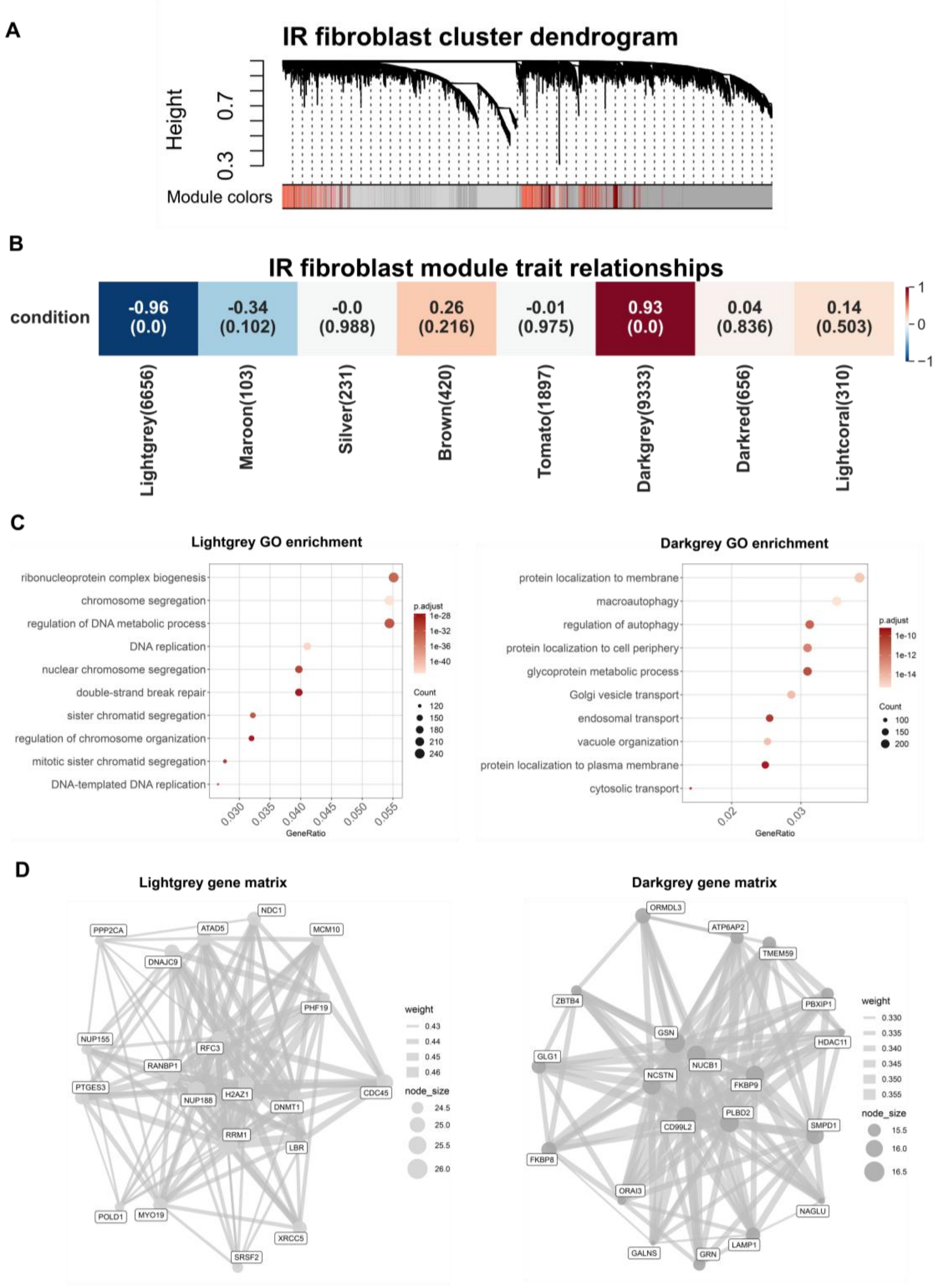

(A) Hierarchical clustering dendrogram of DEGs based on dissimilarity of the topological overlap from the WGCNA. The color bar below the dendrogram indicates module assignments, with each color representing a distinct co-expression module of highly correlated genes.

(B) WGCNA module-trait heatmap relationships from Fig. 3 RNA-seq input of 6 healthy fibroblasts after IR (donors: A18, A27, A31, A37, A48, ID97). Rows represent IR condition, and columns represent module eigengenes. Each cell includes corresponding correlation to the specific module and correlation is indicated by the value in the box. Correlation ranges from -1 to 1 and blue indicates a negative correlation while red indicates a positive correlation. Correlation values are above the number in parenthesis which indicates the p-value of each module's connectivity.

(C) Top 10 GO enrichment terms of genes from the Lightgrey (left) and Darkgrey (right) modules.

(D) Module gene network visualization for the Lightgrey (left) and Darkgrey (right) modules. Gene matrices were made using the BioMart R package. Nodes represent the top module genes ranked by connectivity z-score, node size scales with intramodular connectivity, and edges represent weighted topological overlap matrix (TOM) connections (TOM ≥ 0.001; top 8 connections retained per gene).

Supplemental Figure 9. Per donor graphs from IR iN and IR fibroblasts immunocytochemistry staining of p21, p16, LMNB1, and HMGB1.

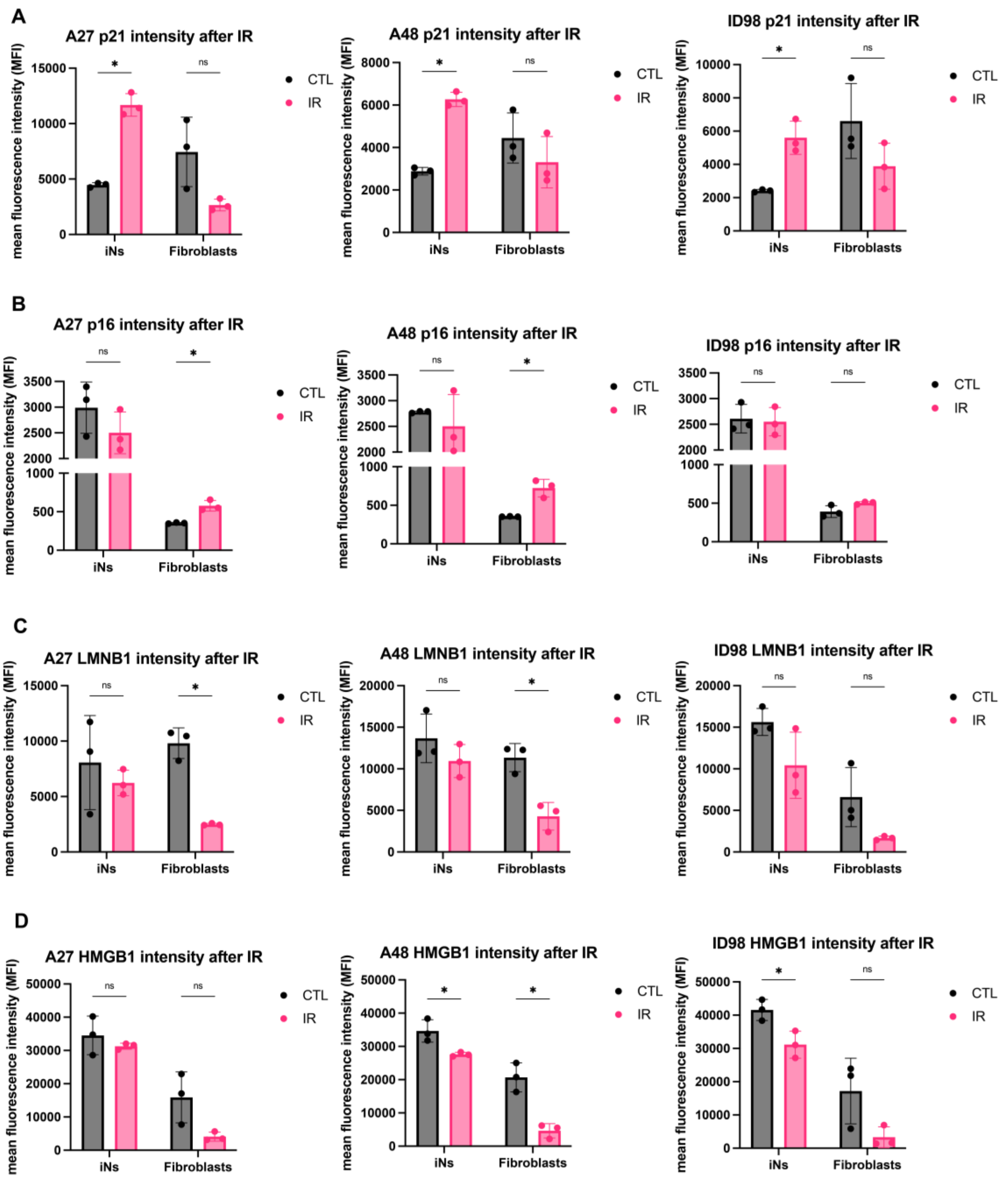

(A-D) IR iNs and IR fibroblasts were stained for p21 (A), p16 (B), LMNB1 (C), and HMGB1 (D) and results from Fig. 6 per donor (donors: A27, A48, ID98; 3 technical replicates per donor in each condition). 0.2% FBS low serum (LS) treatment was applied to CTL and IR fibroblasts to induce quiescence 72 hours prior to collection. All iNs imaged were quantified with automatic MAP2 tracing to measure p21, p16, LMNB1, and HMGB1 only in neuronal nuclei (Supp. Fig. 5). All images were taken on the Harmony Operetta CLS system at 63X unless otherwise stated. Nuclear intensity quantification of images performed by the Harmony analysis system. Statistics were calculated using multiple unpaired t-tests using Welch's correction with Holm–Šídák adjustment for multiple testing with alpha = 0.05. *$p < 0.05$, **$p < 0.01$, ***$p < 0.001$, ****$p < 0.0001$, ns: not significant, nd: not detected.

Supplemental Figure 10. Per donor graphs from IR iN and IR fibroblasts immunocytochemistry staining of γH2AX, 53BP1, and nuclear area.

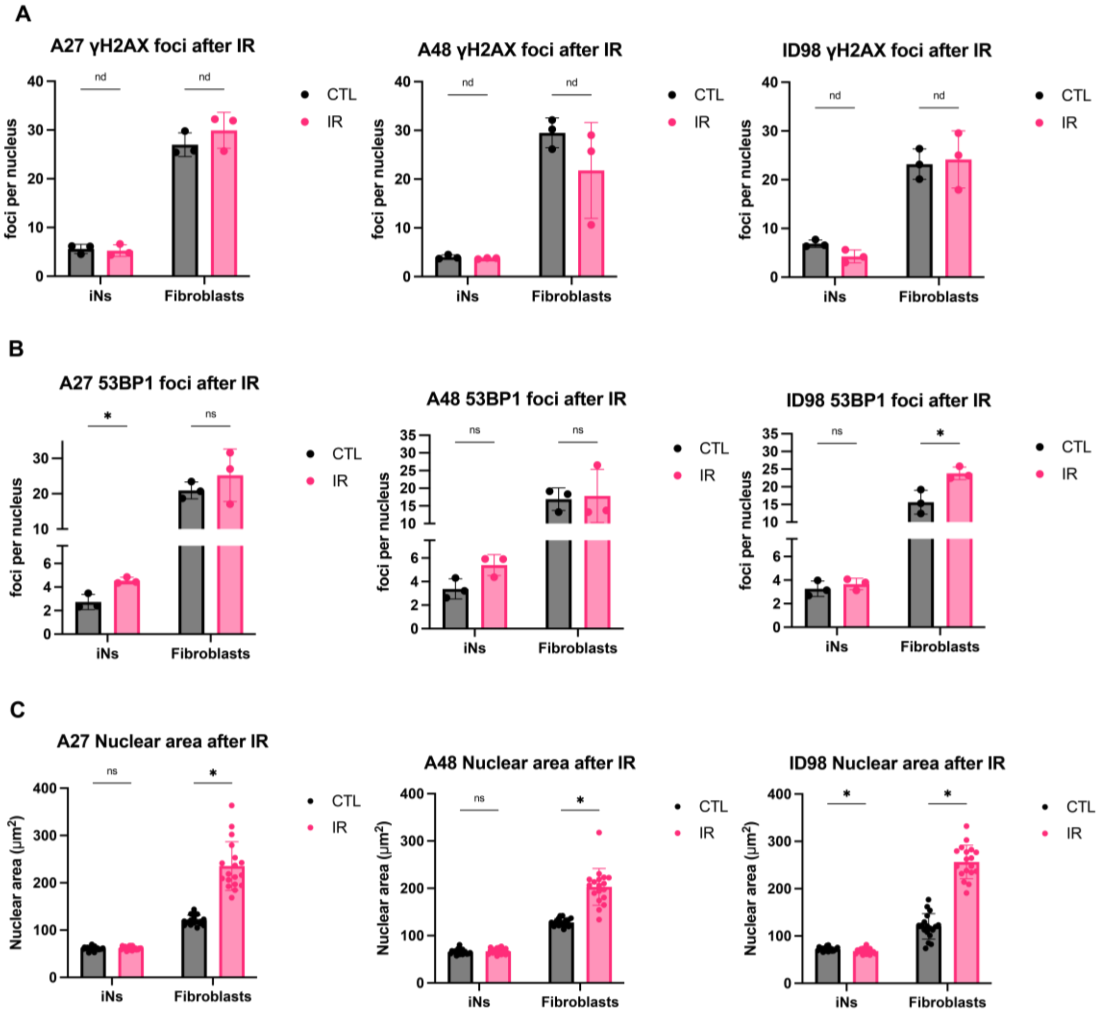

(A-C) IR iNs and IR fibroblasts stained for γH2AX (A), 53BP1 (B), and DAPI (C) and results from Supp. Fig. 3 per donor (donors: A27, A48, ID98; 3 technical replicates per donor in each condition). All iNs imaged were quantified with automatic MAP2 tracing to measure γH2AX, 53BP1, and DAPI only in neuronal nuclei (Supp. Fig. 5). All images were taken on the Harmony Operetta CLS system at 63X. Intensity and foci quantification of images was performed by the Harmony analysis system. Statistics were calculated using multiple unpaired t-tests using Welch's correction with Holm–Šídák adjustment for multiple testing with alpha = 0.05. *$p < 0.05$, **$p < 0.01$, ***$p < 0.001$, ****$p < 0.0001$, ns: not significant, nd: not detected.
